# Genome-wide identification, expression and bioinformatic analyses of GRAS transcription factor genes in rice

**DOI:** 10.1101/2021.06.28.449579

**Authors:** Mouboni Dutta, Anusree Saha, Mazahar Moin, P.B. Kirti

## Abstract

Our group has previously identified the activation tagging of a GRAS transcription factor (TF)gene in the gain-of-function mutant population of rice (*indica* rice variety BPT 5204) screened for water use efficiency (Moin et al, 2016a). This family of GRAS transcription factors has been well known for their diverse roles in gibberellin signaling, light responses, root development, gametogenesis etc. Recent studies indicated their role in biotic and abiotic responses as well. Although this family of TFs received significant attention, not many genes were identified specifically for their roles in mediating stress tolerance in rice. Only *OsGRAS23* (here named as *OsGRAS22*) was reported to code for a TF that induces drought tolerance in rice. In the present study, we have analyzed the expression patterns of rice GRAS TF genes under abiotic (NaCl and ABA treatments) and biotic (leaf samples infected with pathogens, *Xanthomonas oryzae* pv. *oryzae* that causes bacterial leaf blight and *Rhizoctonia solani* that causes sheath blight) stress conditions. In addition, their expression patterns were also analyzed in thirteen different developmental stages. We studied their spatio-temporal regulation and correlated them with *in-silico* studies. Fully annotated genomic sequences available in rice database have enabled us to study the protein properties, ligand interactions, domain analysis and presence of *cis*-regulatory elements in a bioinformatics analysis. Most of the genes were induced immediately after the onset of stress particularly in the roots of ABA treated plants. *OsGRAS39* was found to be very highly expressive gene under sheath blight infection and both abiotic stress treatments while *OsGRAS8*, *OsSHR1* and *OsSLR1* were also responsive. Our earlier functional characterization (Moin et al., 2016a) followed by the genome wide characterization of the GRAS gene family members in the present study clearly show that they are highly appropriate candidate genes for manipulating stress tolerance in rice and other crop plants.

## 1. Introduction

Identification and analysis of transcription factors (TFs) is an essential aspect of functional genomics research. TFs bind to DNA or protein sequences and regulate gene expression (Zhang et al., 2018). They play important roles in almost all cellular functions like growth, development, metabolism, signal transduction, resistance/ tolerance to abiotic and biotic stress factors among others. About 320k TFs from 165 different plant species have been reported. Among them, some important transcription factors include WRKY, MBS, MADS, ARF, AP2/EREBP, HB, SBP, bZIP, GRAS etc. (Zhang et al., 2018; Zijie et al. 2019; Navjot et al., 2020) GRAS group of transcription factors are plant specific proteins, first observed in bacteria and assigned to the Rossman fold methyl transferase superfamily (Zhang et al. 2012). Later, this group radiated towards the ancestors of bryophytes, lycophytes and other higher plants. (Cenci et al., 2017; Zhang et al., 2018). A large number of GRAS genes have been identified in various plant species including 34 in Arabidopsis, 60 in rice, 86 in maize, 106 in *Populus trichocarpa* and many others (Tian et al., 2004; Liu et al., 2014; Guo et al., 2017). The higher number of genes in this gene family indicates that the expansion of the gene family might have happened via segmental and tandem duplication events in evolution and retention of multiple copies post duplication events (Tian et al., 2004; Huang et al., 2015). Till date, the GRAS family of TFs have been studied in 30 different plant species including Arabidopsis, rice, mustard, lotus, tomato, castor bean, poplar, pine, grapevine and others (Cenci et al., 2017). This gene family has been divided into eight subfamilies in Arabidopsis and rice, while the number varied from eight to thirteen in tomato, poplar and castor beans (Tian et al., 2004; Liu et al., 2014; Huang et al., 2015; Wei et al., 2016).

GRAS proteins consist of 400-770 amino acid residues and derive the name from the first three identified members of this family viz. Gibberellin-Acid Insensitive (GAI), Repressor of GAI (RGA) and Scarecrow (SCR) (Pysh et al., 1999; Bolle et al., 2004; Zhang et al., 2017). These genes have a conserved C-terminal region, which forms the GRAS domain and a variable N-terminal region. The conserved region or the GRAS domain comprises five motifs in the following order; leucine heptad repeat I (LHR I), VHIID motif, leucine heptad repeat II (LHR II), PFYRE motif and the SAW motif. (Pysh et al., 1999). The conserved C-terminal domain is responsible for the transcriptional regulation of the genes that exist under their control. The LHR region is required for protein dimerization and the VHIID is necessary for protein-DNA interactions. PFYRE and SAW are the other important regulatory domains that are present in GRAS TFs. Mostly GRAS genes are nuclear localized except PAT1, which is found in the cytoplasm. (Pysh et al., 1999; Tian et al., 2004). The variable N-terminal region consists of intrinsically disordered regions (IDRs,) which are important for molecular recognition during plant development. Due to these IDRs, the GRAS transcription factors are functionally polymorphic (Sun et al., 2012). This gene family integrates environmental and growth regulatory cues and play significant roles in plant development. This family of genes is responsible for a variety of biological functions including gibberellic acid signaling (GAI and RGA of DELLA subfamily and SLR1 of rice) (Pysh et al., 1999; Liu et al., 2014; Vinh et al., 2020), SHR and SCR genes for radial root patterning (Helaritutta et al., 2000), SCL3 for root elongation (Huang et al., 2015), HAM for shoot meristem formation (Stuurman et al., 2002), PAT genes for phytochrome signaling (Bolle et al., 2000), NSP1 and NSP2 for nodulation signaling pathway (Huang et al., 2015) and some others for abiotic and biotic stress responses (Sun et al., 2012; Zhang et al., 2018; Zeng et al., 2019). In many higher angiosperms, several GRAS genes like *ZmSCL7, AtRGA, AtGAI* were shown to have roles in salt stress tolerance in maize and Arabidopsis (Zeng et al., 2019). *PeSCL7* from *Populus* is associated with the modulation of drought and salt tolerance (Ma et al., 2010). *OsGRAS23* (here named as *OsGRAS22*) was shown to induce drought stress tolerance in rice (Xu et al., 2015).

In our previous study (Moin et al., 2016a), we have generated a pool of gain-of-function mutants via activation tagging using tetrameric 35S enhancers and screening of some of these mutants for water use efficiency led to the identification of several genes that were associated with the target trait, the water use efficiency. These interesting gain of function mutants included RNA and DNA helicases (*SEN1* and *XPB2*) (Dutta et al., 2021), and genes for ribosome biogenesis (*RPL6* and *RPL23A*), protein ubiquitination (*cullin4*) and transcription factors like *WRKY 96* and *GRAS* (LOC_Os03g40080) (Moin et al, 2016a). A *GRAS* gene was tagged in the mutant DEB.86 rice line, which showed a high quantum efficiency of 0.82 and a low Δ^13^C value of 18.06‰. Since high photosynthetic efficiency and low carbon isotope ratio are proxies for high water use efficiency, DEB.86 was further analyzed for other phenotypic characters. The activation tagged line DEB.86 exhibited improved plant height with increased tillering and seed yield and had the *ΨOsGRAS4* gene tagged for activation tagging (Moin et al., 2016a).

A total of 60 *GRAS* genes have already been identified in rice (Liu et al., 2014), out of which, *OsGRAS23* has been reported to enhance tolerance to drought (Xu et al., 2015) and *ΨOsGRAS4* has been identified to be associated with enhanced photosynthetic efficiency and water use efficiency with enhanced agronomic features (Moin et al., 2016a). These reports led us to the idea of studying the genome-wide expression analysis of this gene family.

In this study, we have shortlisted forty genes, one gene representing each paralogous group, and provided an experimental basis to identify the potential GRAS genes capable of imparting stress tolerance in rice. We have analyzed the genes selected in the GRAS family for their spatio-temporal and stress induced expression. The phylogenetic relationship among GRAS proteins, their genetic arrangements and structure, *in-silico* analysis of putative promoter elements and protein properties were also studied. This study helps in the identification of important GRAS genes for stress tolerance, which aids in their further functional characterization.

## 2. Materials and methods

### 2.1 Retrieval and nomenclature of GRAS sequences

Our previous work on gain of function mutants generated through activation tagging technology using the tetrameric 35S elements identified a GRAS gene as a potential player in enhancing water use efficiency in rice (Moin et al., 2016a). Also, Xu et al. (2015) suggested that *OsGRAS23* is involved in inducing drought stress responses in rice. This has led us to undertake literature search in the present study and we observed that Tian et al. (2004) have identified 57 GRAS genes in rice. We searched the accession numbers of all 57 genes in NCBI and did a BLASTN search in rice genome database (RGAP-DB, Orygenes DB), and retrieved the locus numbers of 47 genes. Simultaneously, we did a key word search of GRAS, DELLA, Scarecrow, Monoculm, Chitin-inducible gibberellin-responsive protein, Gibberellin response modulator protein, Nodulation signaling pathway and Short Root, and combined the search results with the 47 genes retrieved from the literature search. We matched our list of 60 genes with that of Cenci and Rouard (2017) and followed the same nomenclature. For more clarity, we had performed a protein database search for the GRAS domain in NCBI, SMART, Prosite and Pfam databases. We had selected 40 genes, one each from all the paralogous groups for our analyses.

### 2.2. Genomic distribution of GRAS genes

The coordinates of all 60 GRAS genes were obtained from RGAB-DB and were fed in the NCBI Genome Decoration Page. The outputs were combined and the genes were marked for understanding the genomic distribution of *OsGRAS* genes.

### 2.3. Phylogenetic relationship of rice GRAS genes

In order to understand the evolutionary relationships between the rice GRAS genes, we aligned the amino acid sequences in MEGA7 software followed by the construction of an unrooted phylogenetic tree. The tree was constructed using the Neighbour Joining method with a bootstrap value of 1000.

### 2.4. Motif arrangements and organization of GRAS genes

All the 40 GRAS genes were subjected to MEME suite for conserved motif analysis using default parameters. The number of motif scan was set to 10. Based on the previous article of Pysh et al. (1999), the MEME-motifs were further classified into conserved GRAS motifs. The gene organization was studied by subjecting the genomic and coding sequences in the Gene Structure Display Server (GSDSv2). The number of exons, introns, untranslated regions (UTRs) etc. were noted.

### 2.5. *In-silico* analysis of the putative promoter region

The *cis*-acting elements in the promoter regions play a major role in the coordinated expression of the genes. Hence, it is crucial to identify these regulatory elements in order to correlate the expression data with the genetic components. We retrieved ≤1kb upstream sequences of all 40 selected GRAS genes under study from the rice genome database and identified important elements responsible for biotic and abiotic stress responses in them. The identification of the elements was performed by subjecting the sequences in PlantCARE (Cis-Acting Regulatory Elements) database and manually mapping them on the chromosomes.

### 2.6. Biochemical properties of GRAS proteins

The sequences of forty GRAS genes that were shortlisted were subjected to ExPASyProtParam tool to gauge their encoded proteins with amino acid length, molecular weight and theoretical isoelectric points (*p*I). The three-dimensional structures of the proteins and their ligand interactions were studied using 3DLigandSite software (Wass et al., 2010). The structures were then subjected to Phyre2 (Protein Homology/Analogy Recognition Engine v2; Kelley et al., 2015) program for analysis of the protein secondary structure composition. This tool gives an idea of the percentage of secondary structures in a protein i.e. the percentage of α-helix, β-sheets and disordered regions in proteins. The SMART (Simple Modular Architecture Research Tool) online tool was used to analyse the protein domains and their low complexity regions (LCRs). ExpasyProtParam tool also indicated the GRAVY indices of the proteins, which provide information regarding the hydrophobicity of proteins. The localization and existence of transmembrane helices of the genes were predicted using TargetP-2.0 and TMHMM software, respectively.

### 2.7. Preparation of Plant material for studying the gene expression under native and stress conditions

For simulated abiotic stress experiments, BPT-5204 (Samba Mahsuri) rice seeds were surface sterilized using 70% ethanol for 50 sec followed by 4% aqueous sodium hypochlorite solution for 15 min and five washes with sterile double distilled water, each of one minute duration. The sterile seeds were grown on Murashige and Skoog medium for 7 d under a 28 ± 2°C for 16 h/ 8 h photoperiodic cycle (Saha et al., 2017). The seedlings were then subjected to NaCl (250 µM) and ABA (100 µM) stress conditions for 60 h. Shoot and root samples were collected periodically at 0 h, 15 min, 3 h, 12 h, 24 h and 60 h after the onset of stress. The untreated samples were taken as controls for normalization of gene expression.

For studying the native expression patterns of the GRAS genes, tissue samples from thirteen regions in rice seedlings were collected (Moin et al., 2016b, Saha et al., 2017). These included embryo and endosperm from 16 h soaked seeds, plumule and radicle from 3 d old germinating seeds, shoot and root tissues from 7 d old seedlings and shoot, root, root-shoot transition region, flower, spikes and grain samples from mature 20 d old plants post-transfer to the greenhouse.

In order to study the expression of GRAS genes under biotic stress conditions, leaf samples of one month old rice plants infected with *Xanthomonas oryzae* pv. *oryzae* (*Xoo* that causes Bacterial Leaf Blight, BLB) and *Rhizoctonia solani* (that causes Sheath Blight, SB) were taken post 20 d and 25 d of infection, respectively. Samples from plants of the same age without the pathogen treatment were taken as the controls. The infection protocol was followed as per Saha et al. (2017).

### 2.8. c-DNA preparation and Quantitative Real-Time PCR (qRT-PCR)

The plant material collected was used to isolate of RNA using Tri-reagent following manufacturer’s protocol (Takara Bio, UK) and c-DNA was prepared using 2 µg total RNA samples (Takara Bio, UK). The c-DNA samples were diluted ten times and an aliquot of 2 µl of each sample per reaction was used for qRT-PCR. All the primers were designed using Primer3 software and 10 µM primer concentration was used per reaction. The PCR program included an initial denaturation step of 94°C for 2 min followed by 40 cycles of second denaturation of 30 sec, annealing for 25 sec and extension at 72°C for 30 sec. The samples for the current study were taken in biological and technical triplicates and the fold changes were calculated using the ΔΔC_T_ method (Livak and Schmittgen, 2001). Rice *actin* and *β-tubulin* genes were used as two housekeeping genes for internal normalization. For abiotic and biotic expression studies, housekeeping genes and individual control samples were used for double normalization. In contrast, single normalization was performed using the C_T_ value of housekeeping genes for native expression studies. The graphs were generated using MORPHEUS program and GraphPad Prism software. One way ANOVA was performed using SigmaPlot software for discerning the significance of statistical differences between samples.

## 3. Results

### 3.1. Chromosomal distribution of GRAS genes in rice genome

Liu and Widmer (2014) showed that there are 60 GRAS genes in the genome that are distributed on 10 out of 12 chromosomes of rice. Based on the literature and database search, we observed that chromosome 8 and 9 did not carry any GRAS genes. The number of genes on a single chromosome ranged from a minimum of two on chromosome 10 to a maximum of twelve on chromosome 11. Among the rest, a total of nine genes were located on chromosome 3, while chromosome 1, 7 and 12 carried six genes each, chromosome 2, 4 and 5 exhibited five genes each and chromosome 6 had four genes (Fig. 1). Out of the 60 genes located, we have shortlisted 40 genes for our study with one representative from each paralogous group selected.

**Figure 1:**
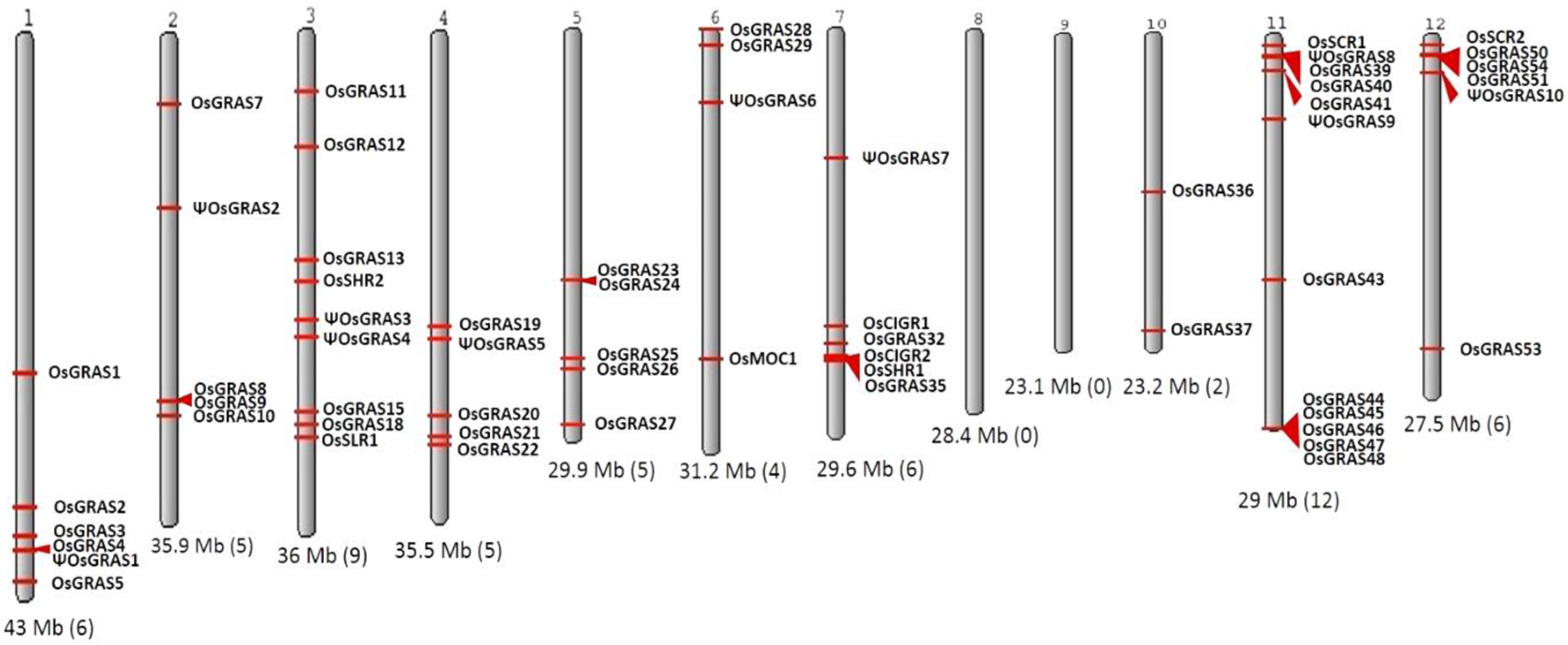
Chromosomal distribution of GRAS genes in rice. Karyotypic representation of rice chromosomes obtained from NCBI Genome Decoration Page. Rice genome carries 60 GRAS genes, which are represented in the figure with red arrows indicating the position of each gene. The size of each chromosome and the number of genes present are provided below each in each bracket.

### 3.2. Analysis of evolutionary relationship of *OsGRAS* genes

In order to understand the evolutionary relationship among the rice GRAS family of genes, we subjected the retrieved sequences to a phylogenetic analysis (Fig. 2) in MEGA7 software. A total of 16 different clusters were observed. These clusters were divided into 14 subfamilies based on a previous report of Cenci and Rouard (2017). Members belonging to the same subfamily were found to cluster together except DLT and PAT subfamilies where some genes belonging to different orthologous groups (according to Cenci and Rouard, 2017) formed separate clusters. Each cluster has been colour coded in the figure. The number of genes found in each subfamily included four in SCL3, three each in SCR, NSP2 and HAM, one in RAM, LS, SCL4/7and SCLA, two in DELLA, DLT, SHR and SCL32, six in PAT and nine in LISCL. LISCL was found to be the largest subfamily with maximum member of genes getting clustered. *ΨOsGRAS4* and *ΨOsGRAS9* were placed close to LISCL family since these sequences were still unclassified. The highly expressed genes under biotic and abiotic stress conditions belonged to SCL3, SHR, DELLA, HAM and PAT subfamilies.

**Figure 2:**
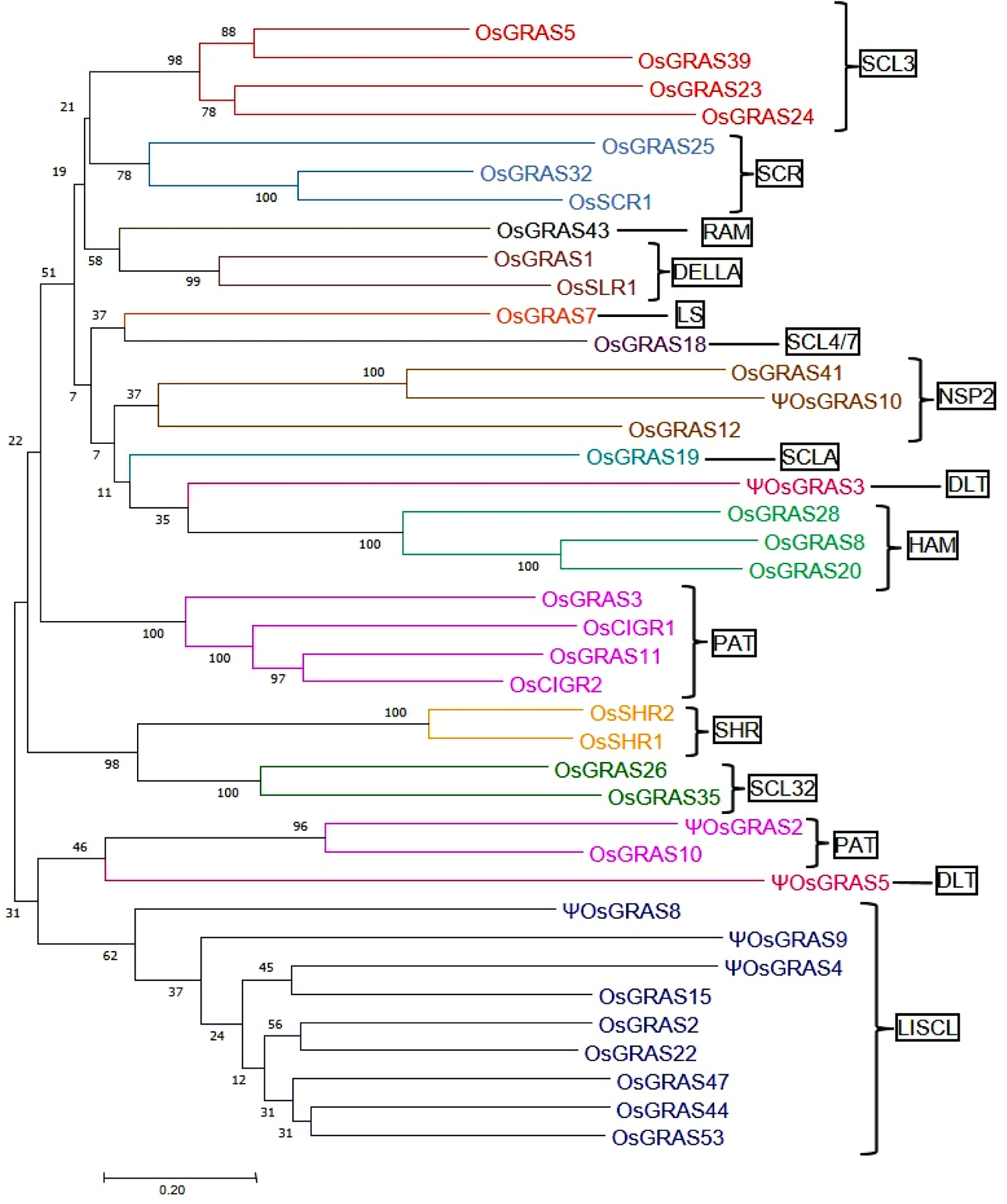
Phylogenetic analysis of *OsGRAS* genes. An unrooted phylogenetic tree showing the evolutionary relationship of *OsGRAS* genes. The tree was constructed using the Neighbour Joining method in MEGA7 software with a bootstrap value of 1000. The number at each node represents the percentage bootstrap values. Based on the previous literature, the genes have been divided into 14 subfamilies (mentioned in boxes) and each subfamily has been colour coded.

### 3.3. Analysis of GRAS motifs and gene organization

The amino acid sequences of selected 40 genes were subjected to MEME analysis for identifying the conserved motifs in rice GRAS gene encoded proteins. A total of ten motifs were identified, which corresponded to LHR I (motif 5, 9), VHIID (motif 2, 3, 10), LHR II (motif 8), PFYRE (motif 4, 7) and SAW (motif 1, 6) motifs (Fig. 3 and Fig S1). The C-terminal domain was found to contain the conserved GRAS motifs as reported earlier in literature. However, not all genes exhibited all the ten MEME-motifs. PAT and LISCL subfamilies carried all the ten domains, while others like SCR lacked motif 1. Proteins belonging to same subfamily had similar motif composition.

**Figure 3:**
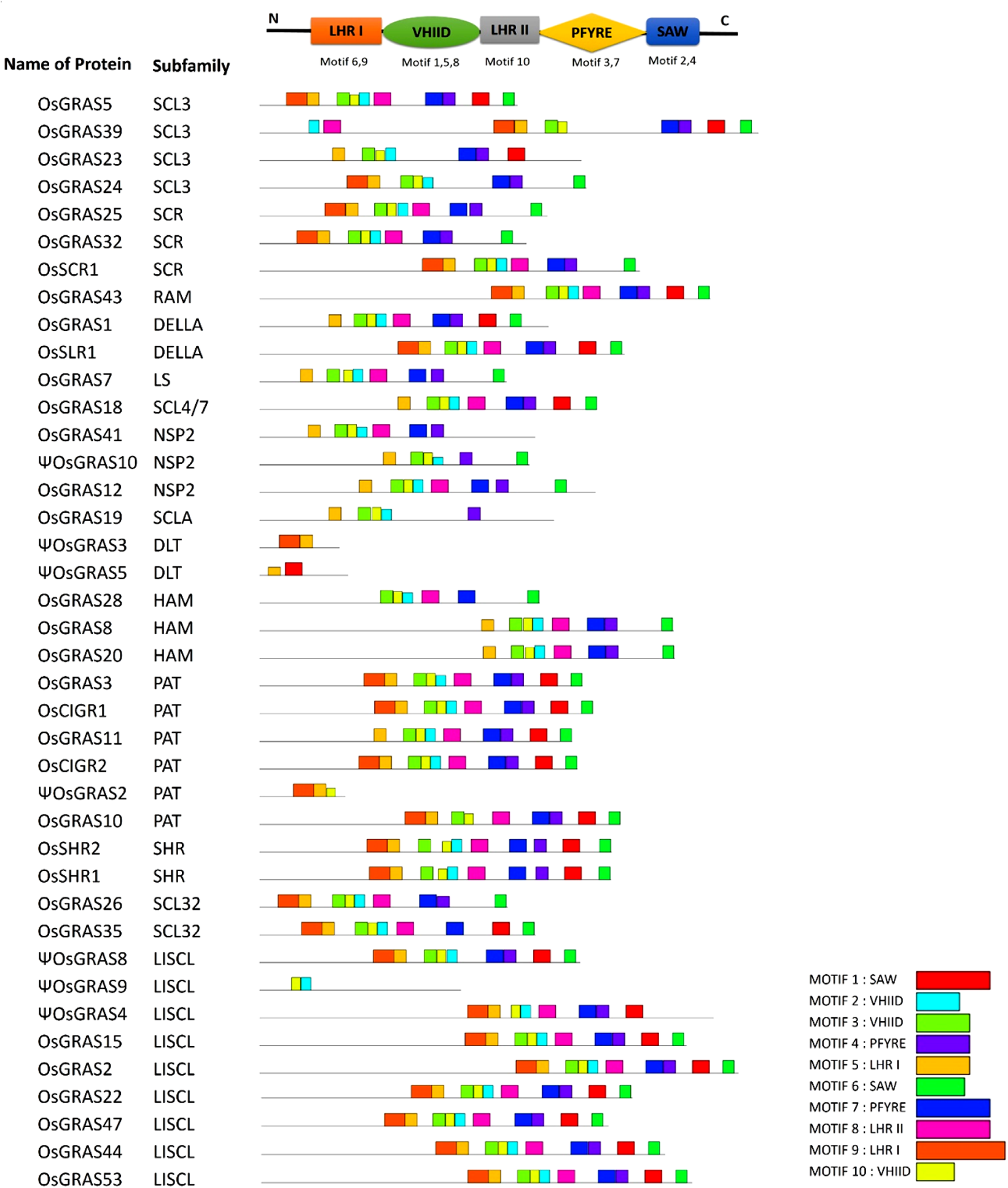
MEME-motif analysis of *OsGRAS* genes. Figure showing the identified MEME-motifs of *OsGRAS* genes. The conserved GRAS-motifs are provided at the top. A search for 10 MEME-motifs was done and each of them has been assigned to the corresponding GRAS-motifs. Each coloured box represents one motif and the legend has been provided below. The genes were organized based on their subfamilies.

The genomic and cDNA sequences of all the selected 40 genes were subjected to GSDS server to observe the organization of different GRAS genes selected from each of the paralogous groups (Fig S2). Based on the map that was generated by the server, it was observed that the genes varied in length and the distribution of exons, introns and untranslated regions (UTRs). The majority of genes (31 out of 40 genes studied) lacked introns in their gene structure and were only composed of exonic sequences and UTRs. *OsGRAS11* exon is flanked by a long stretch of UTR at its 5’ and 3’ ends. It completely lacked introns and is the longest gene in this study (6.7Kb). Ten genes were observed to contain only coding sequences in their structure without any introns and UTRs. Among them, *ΨOsGRAS3* had the smallest sequence of only 414 bp. Nine genes carrying introns only in their structure were *OsGRAS3, OsGRAS39, OsGRAS41, OsGRAS43, OsSCR1, ΨOSGRAS4, ΨOsGRAS8, ΨOsGRAS9* and *ΨOsGRAS10.* The number of intronic sequences among the genes varied between one (*OsSCR1* and *ΨOsGRAS10*) to a maximum of seven (*ΨOsGRAS4*). All of them showed low (*OsGRAS43*), moderate (*OsGRAS3*, *ΨOSGRAS4* and *ΨOSGRAS8*) and very high (*OsGRAS39*, *OsGRAS41*, *OsSCR1*, *ΨOsGRAS9* and *ΨOsGRAS10*) expression levels under abiotic and biotic stress conditions. Six out of nine genes (*OsGRAS41, OsGRAS43, ΨOSGRAS4, ΨOsGRAS8, ΨOsGRAS9* and *ΨOsGRAS10)* did not exhibit any UTRs in their structure and were solely composed of introns and exons. The details of genetic organization of rice GRAS genes have been provided in the Supplementary Table S1.

### 3.4. Putative promoter analysis of GRAS genes and the search for *cis*-regulatory elements

Since diverse expression patterns were observed for diffenernet GRAS genes under abiotic and biotic stress conditions, we tried to correlate their expression patterns with the putative regulatory sequences observed in their upstream regions. In order to achieve this correlation, we retrieved 1 Kb sequences from 5’ upstream region of each gene under study from the rice genome database and subjected them to an *in-silico* analysis for the identification of the *cis*-putative regulatory elements observed in them. A total of eighteen stress responsive elements were observed in the upstream putative promoter region of the GRAS genes. These included ABRE or ABA responsive elements, CCAAT box and MYB sites for binding of MYB transcription factors responsive to drought inducibility, binding site for MYC transcription factors for defence responses, DRE or dehydration responsive elements, STRE or stress responsive elements, TC-rich repeats for defence and stress responses, and the LTR or low temperature responsive element. Several phytohormones and wound responsive elements were also observed in their upstream regions, which included TCA-element for salicylic acid responses, CGTCA-motif or TGACG-motif as a methyl jasmonate responsive element, GARE-motif, TATC-box and P-box for gibberellin responses, ERE as ethylene responsive elements, TGA-element or AuxRR core or AuxRE for auxin responses, WUN-motif and WRE for responses against wounding, box-S for wounding and pathogen elicitation, and the W-box for binding of WRKY transcription factors.

*OsGRAS39*, the highly expressive gene under both biotic and abiotic stress conditions in the present study had three copies each of MYB binding factor sites and CGTCA-motif, five copies of STRE, two copies of ABRE and one copy each of DRE, TC-rich repeats and CCAAT-box justifying its expression under different stress treatments. Other responsive genes in both the stresses like *OsGRAS8*, *OsSHR1* and *OsSLR1* had combinations of MYB, STRE, ERE, WUN, TCA, CGTCA and MYC elements in their putative promoter regions. Apart from these, *OsGRAS8* exhibited ABRE, LTR and W-box elements, *OsSHR1* carried a DRE element and *OsSLR1* had copies of TATC, WRE and TC-rich elements. *ΨOsGRAS5*, only expressive gene in the shoot region had two copies each of MYB and MYC binding elements and three copies of ABRE. Other important abiotic stress responsive genes like *ΨOsGRAS2* and *OsSCR1* were observed to have multiple copies (upto six) of ABRE, MYB and MYC elements, STRE elements and ERE, CGTCA, GARE and WRE motifs in their 5’ upstream regions. *OsCIGR1* that was found to be highly induced under biotic stress conditions carried ten copies of ABRE, seven copies of STRE, five copies of CGTCA element and one copy each of CCAAT-box, DRE, MYB, MYC and WRE. Other expressive genes under biotic stress conditions included *OsGRAS2*, *ΨOsGRAS3, OsGRAS19, OsGRAS20* and *OsGRAS23,* which had combinations of TCA-elements, W-box, WRE, ERE, AuxRE, CGTCA-box, box-S and WUN elements apart from other stress responsive elements. The function of each elements has been provided in Table 1 and the physical mapping of the important stress responsive elements on the putative promoter regions of the genes was provided in Fig 4.

**Figure 4:**
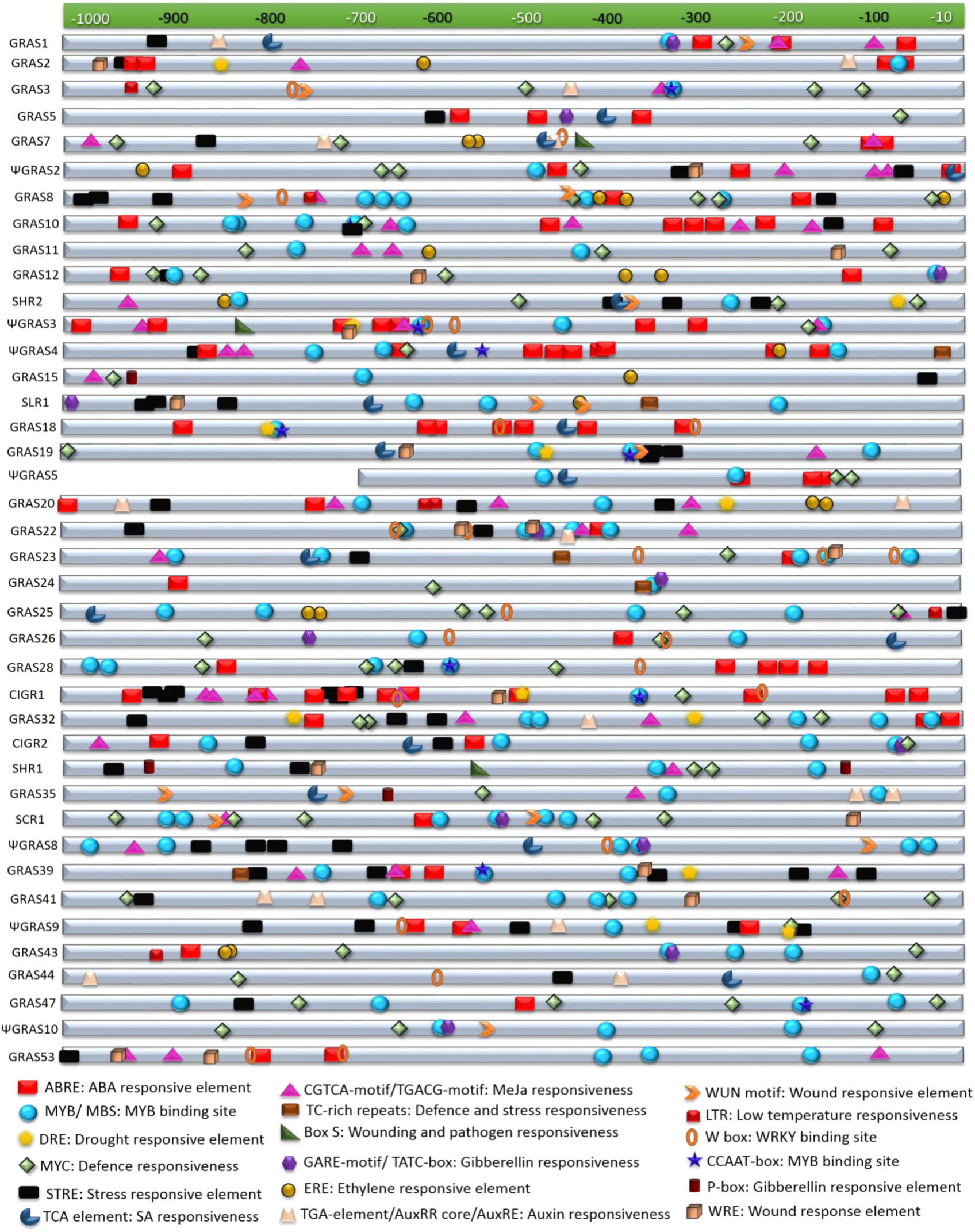
*In-silico* analysis of putative promoter regions of GRAS genes. The selected GRAS genes were subjected to *in silico* analysis for *cis*-regulatory elements in their putative promoter regions (sequence retrieved from about ≤1kb upstream region). This was performed in PlantCARE database and the figure was prepared by mapping the stress regulatory elements in the each of the sequences. The index for each element along with its functions are mentioned below the figure.

**Table 1:** List of cis-regulatory elements and their functions

| Name of *cis*-element | Function |
| --- | --- |
| ABRE | ABA responsive element (Choi et al. 2000) |
| MYB/MBS | MYB binding site for drought inducibility (Ambawat et al. 2013) |
| DRE | Dehydration responsive element (Narusaka et al. 2003) |
| MYC | Transcription factor for stress responses, helps in dehydration induced expression of genes (Tran et al. 2004) |
| STRE | Stress responsive element (Hwang et al. 2010) |
| TCA element | Element for salicylic acid responsiveness (Wei et al. 2013) |
| CGTCA-motif/TGACG-motif | Methyl-Jasmonate responsive element (Wang et al. 2011) |
| TC-rich motifs | Responsible for defense and stress, transcription regulation (Bernard et al. 2010; Liu et al. 2017) |
| Box S | Responsive to wounding and pathogen elicitation (Yin et al. 2017); Stress responsiveness (Ding et al. 2019) |
| GARE-motif/TATC-box | Gibberellin responsive element (Bastian et al. 2010) |
| ERE | Element for ethylene responses (Sanchez and Singh 2002) |
| TGA-element/AuxRR core/ AuxRE | Element for auxin response (Sakai et al. 1996) |
| WUN motif | Wound responsive element for biotic stress (Xu et al. 2011) |
| LTR | Low temperature responsive element (Zhang et al. 2020) |
| W box | Binding sites for WRKY transcription factors (Dhatterwal et al. 2019) |
| CCAAT box | Binding site for MYB transcription factors |
| P-box | Gibberellin responsiveness (Zhang et al. 2020) |
| WRE | Wound responsive element (Whitbred and Schuler 2000) |

### 3.5 Properties of GRAS proteins, their ligand interactions and domain analysis

We studied the properties of 40 shortlisted GRAS proteins like amino acid length (aa), molecular weight (KDa) and theoretical *p*I through ExPASyProtParam program. It was observed that the proteins had a molecular weight ranging from 15 kDa (ΨOsGRAS3) to 94 kDa (OsGRAS39). ΨOsGRAS3 showed a minimum amino acid (aa) length of 137 aa while OsGRAS39 had a maximum length of 854 aa. The *p*I of the proteins ranged from acidic to basic (4.5-10.1) with only eight proteins having a *p*I of more than 7. The majority of the proteins fall under the *p*I range of 4-7. Likewise, the remaining 32 proteins were found to be in the acidic range i.e. *p*I <7. This is because of the observation that the proteins carried more negatively charged (acidic) amino acid residues like Aspartic acid and Glutamic acids in their composition as compared to basic amino acid residues. Only OsGRAS39 was found to have an equal number of acidic and basic residues in its composition. According to TargetP-2.0 server, OsGRAS39 was predicted to be localized to the chloroplast while no signal peptides for chloroplast or mitochondria could be specified by the tool for the rest of the proteins.

We have also analyzed the proteins for their three dimensional structures and ligand binding residues in the 3DLigand site and the structures were submitted to the Phyre2 program to analyse their secondary structures like the percentage of disordered regions, α-helix and β-sheets. ΨOsGRAS3 showed a maximum of 71% and OsGRAS8 had a minimum of 31% of α-helical structure. Similarly, maximum (14%) extent of β-sheets were noticed in the secondary structure of OsGRAS32. No β-sheets were present in ΨOsGRAS2 and ΨOsGRAS3. Several metallic and non-metallic ligands were also observed to be interacting with the GRAS proteins, which included Mg^+2^, Ca^+2^, SAM, SAH, NAP, NAD, ATP, Zn^+2^ and Ni^+2^. The three dimensional structures of the proteins along with their interacting ligands have been provided in the Fig. S3.

Low complexity region (LCR) are repetitive amino acid sequences found abundantly in the eukaryotic proteins. These play essential roles in protein-protein and protein-nucleic acid interactions (Toll-Riera et al., 2012). It was noted that the number of LCRs in each of the proteins varied from none to a maximum of eight in OsGRAS20 and OsGRAS43, respectively.

Grand average of hydropathicity index or GRAVY index indicates the hydrophobicity of a protein taking into consideration its charge and the size. Usually GRAVY values range from −2 to +2 with more positive values indicating hydrophobicity and more negative values indicating hydrophilicity (Morel et al., 2006). Seven proteins had a positive GRAVY value while the rest 33 proteins had a values lesser than zero, which indicated that the majority of the GRAS proteins are hydrophilic. The list of all the observations have been provided in the supplementary table S2.

In order to study the domains present in the genes, we utilized the SMART online tool and observed that all the proteins had at least one GRAS domain with ΨOsGRAS4, ΨOsGRAS8, OsGRAS39, ΨOsGRAS10 exhibiting two GRAS domains. Among them, ΨOsGRAS4 and ΨOsGRAS10 had two internal repeats designated as RPT1 along with two GRAS domains. One DELLA domain and one SCOP domain in addition to the GRAS domain were found in OsSLR1 and OsGRAS18, respectively. DELLA proteins are transcriptional regulators, which function in gibberellic acid signaling by binding with GA receptor, GID1 followed by proteasomal degradation of DELLA domain (Murse et al., 2008). OsGRAS41 had a transmembrane region, OsGRAS43 and OsGRAS53 had two RPT1 domains (internal repeats) along with their single GRAS domains. The detailed list of the domains and the LCRs with their sequences have been provided in the Table S3. The presence of transmembrane domain in OsGRAS41 was further confirmed through TMHMM software.

### 3.6. Expression analysis under simulated abiotic stress conditions

We have identified a GRAS transcription factor as a potential stress tolerance gene by screening a pool of gain-of-function mutants in rice in our previous study (Moin et al., 2016a). Another report by Xu et al. (2015) suggested the role of *OsGRAS23* (reported as *OsGRAS22* in this study) in drought tolerance in rice. These observations have prompted us to analyse the differential expression pattern of GRAS family of transcription factors under the influence of biotic and simulated abiotic stress conditions in the present study. We have analyzed the expression patterns of 40 selected genes separately in shoot and root tissues at six different time points for two abiotic (NaCl and ABA) and two biotic (BLB and SB) stresses. The native expression patterns of these genes in 13 different tissues were also studied.

Based on the pattern of expression, we have divided the genes as immediate early (IE), early (E) and late (L) responsive genes. Some genes were expressed up to 100 folds after the incidence of stress. Thus, the genes were also categorized as expressive (2-10 fold), moderately expressive (10-30 fold) and highly expressive (≥30 fold) types. Genes showing an upregulation of ≥2 folds were considered as expressive.

The majority of the genes got upregulated in the root (Fig. 5a, b) compared to the shoot (Fig. 5c, d). As indicated in the pie chart, about 55-60% of the total genes showed IE type expression under both NaCl and ABA treatments. NaCl, however, induced more early (12.5%) responsive genes than late (2.5%) whereas, ABA induced more late genes (12.5%) than early (2.5%). The list of the expressed genes has been provided in Fig. 5. More than half of IE genes continued their expression till 60 h of treatment, while some others became downregulated or showed no expression at all later during the experimental timeline. Under ABA treatment, all highly upregulated genes i.e. *ΨOsGRAS2, OsSHR1, OsSCR1* and *OsGRAS39* were IE type and their expression persisted till the last time point of treatment i.e. 60 h. Other IE type genes showed a split before increasing their expression at subsequent time points. Only *OsGRAS39* was highly expressive under both ABA and NaCl treatments (100 fold and 65 fold, respectively). *OsGRAS2*, *ΨOsGRAS2, OsGRAS25, OsGRAS35* and *OsSCR1* under NaCl and *OsGRAS22* under ABA were early (E) expressed genes respectively, while *ΨOsGRAS9, OsGRAS3, OsGRAS11* and *OsGRAS26* under ABA and *ΨOsGRAS9* became upregulated under NaCl treatment with late (L) expression.

**Figure 5:**
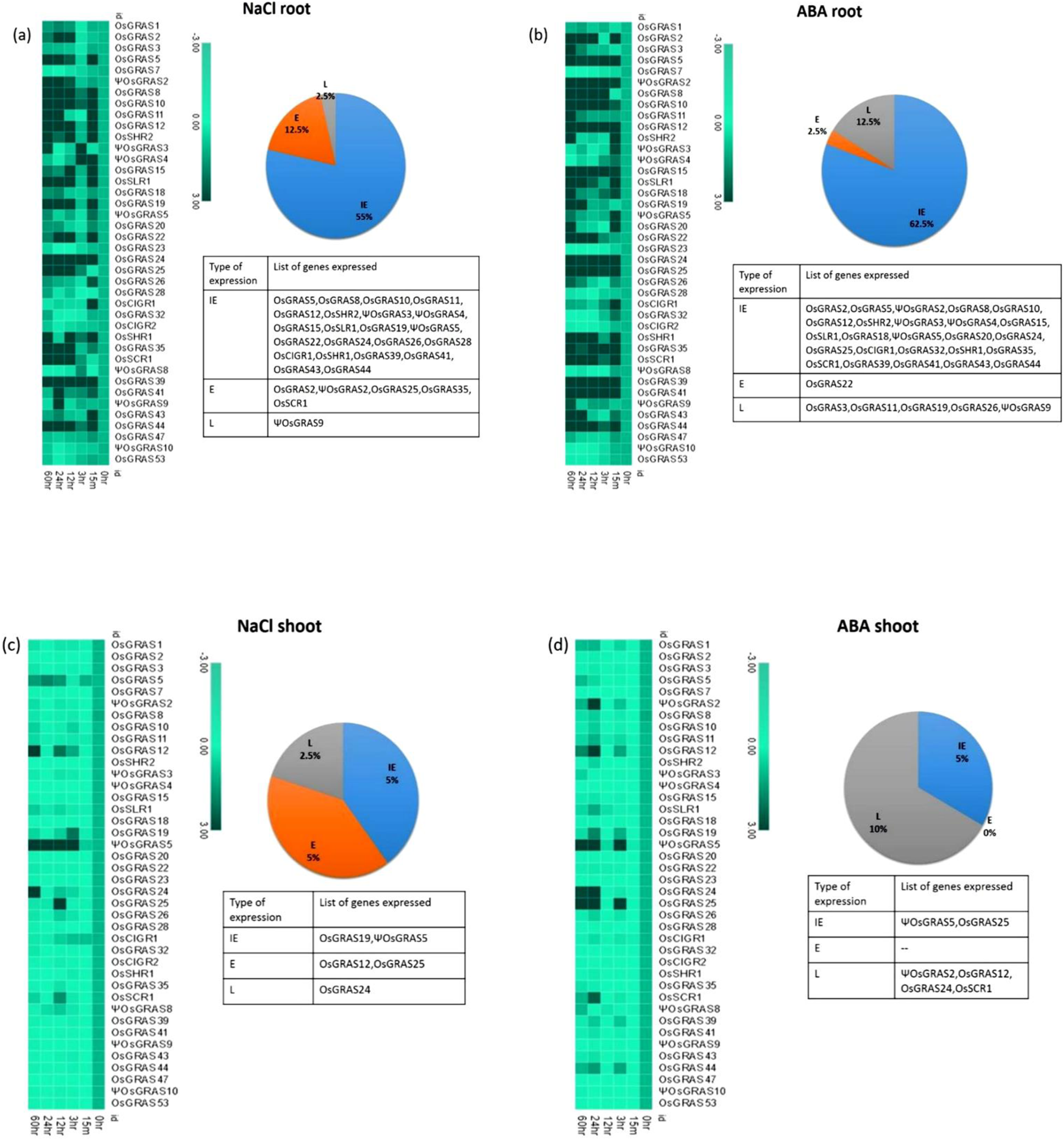
Expression analysis of GRAS genes under abiotic stress. Heat map representation of temporal expression pattern of GRAS genes developed using MORPHEUS program. 7 d old seedlings were subjected to NaCl (250µM) and ABA (100µM) treatments and the obtained quantitative real-time values were double normalized using rice *actin* and *tubulin* as the internal reference genes and that of the unstressed samples using the ΔΔC_T_ method. The experiment was conducted separately for root (4a, b) and shoot (4c, d) tissues. Percentage of genes upregulated under NaCl and ABA treatments is represented in the form of a pie chart beside their corresponding heat maps. The genes were separated based on their time point(s) of expression and annotated as immediate early (IE), early (E) and late (L) expressive genes. The name of the genes is provided in the list below.

Twelve and thirteen genes (30 and 32%) were mild to moderately expressed, respectively under the ABA treatment, whereas seven genes (17%) were moderately expressive under NaCl treatment and rest 19 genes (47%) exhibited mild expression. Nine genes (22%) under ABA and thirteen (32%) genes under NaCl treatment were either downregulated or showed no change in the level of expression. Among them, *OsGRAS7, OsGRAS23, OsGRAS28* and *ΨOsGRAS8* were downregulated under both treatments. *ΨOsGRAS3, ΨOsGRAS4, OsCIGR1* and *OsGRAS32* under ABA treatment and *OsCIGR1* under NaCl treatment showed an immediate expression, but was either downregulated or showed no expression at subsequent time points

Not many genes were expressed in the shoot. However, *ΨOsGRAS5* is the only gene, which showed moderate expression (25-30 fold) under both ABA and NaCl treatments. This gene was IE type maintaining its expression till 60 h under NaCl, but showed a split before reaching a peak under the ABA treatment. On the contrary, it showed low expression (2-3 fold) in root tissues under ABA and NaCl treatments. Among the other genes that were mildly expressive in both root and shoots were *ΨOsGRAS2, OsGRAS12, OsGRAS19, OsGRAS24, OsGRAS25* and *OsSCR1*. The rest of the genes were mainly downregulated or did not show any change in expression in shoot tissues under both the stress treatments. The expression level of all the genes studied has been provided in Table 2.

**Table 2:** List of GRAS genes and their expression pattern under NaCl and ABA

| Gene name | Locus number | Regulation<br>(up/down) | Type of<br>Response | Maximum<br>fold change | Regulation<br>(up/down) | Type of<br>Response | Maximum<br>fold change |
| --- | --- | --- | --- | --- | --- | --- | --- |
|  |  | **Root** |  |  | **Shoot** |  |  |
| **NaCl** |  |  |  |  |  |  |  |
| OsGRAS1 | LOC_Os01g45860 | DOWN | - | - | - | - | - |
| OsGRAS2 | LOC_Os01g62460 | UP | E(2.9) | 3.3 | - | - | - |
| OsGRAS3 | LOC_Os01g65900 | DOWN | - | - | - | - | - |
| OsGRAS5 | LOC_Os01g71970 | UP | IE (5.5) | 6.1 | - | - | - |
| OsGRAS7 | LOC_Os02g10360 | DOWN | - |  | DOWN | - | - |
| ΨOsGRAS2 | LOC_Os02g21685 | UP | E(2.6) | 18.8 | DOWN | - | - |
| OsGRAS8 | LOC_Os02g44360 | UP | IE(5.6) | 12 | DOWN | - | - |
| OsGRAS10 | LOC_Os02g45760 | UP | IE(12.4) | 12.4 | DOWN | - | - |
| OsGRAS11 | LOC_Os03g09280 | UP | IE(3.8) | 4.8 | DOWN | - | - |
| OsGRAS12 | LOC_Os03g15680 | UP | IE(9.5) | 9.5 | UP | E(2.1) | 6.8 |
| OsSHR2 | LOC_Os03g31880 | UP | IE(3.6) | 4.3 | DOWN | - | - |
| ΨOsGRAS3 | LOC_Os03g37900 | UP | IE(3.7) | 3.7 | DOWN | - | - |
| ΨOsGRAS4 | LOC_Os03g40080 | UP | IE(3.0) | 16.2 | DOWN | - | - |
| OsGRAS15 | LOC_Os03g48450 | UP | IE(3.8) | 3.8 | DOWN | - | - |
| OsSLR1 | LOC_Os03g49990 | UP | IE(3.3) | 11 | DOWN | - | - |
| OsGRAS18 | LOC_Os03g51330 | - | - | - | DOWN | - | - |
| OsGRAS19 | LOC_Os04g35250 | UP | IE(4.2) | 6.4 | UP | IE(2.1) | 2.1 |
| ΨOsGRAS5 | LOC_Os04g37440 | UP | IE(2.2) | 2.2 | UP | IE(7.7) | 25.2 |
| OsGRAS20 | LOC_Os04g46860 | DOWN | - | - | DOWN | - | - |
| OsGRAS22 | LOC_Os04g50060 | UP | IE(3.2) | 4.6 | DOWN | - | - |
| OsGRAS23 | LOC_Os05g31380 | DOWN |  |  | DOWN | - | - |
| OsGRAS24 | LOC_Os05g31420 | UP | IE(4.0) | 8.6 | UP | L(4.6) | 4.6 |
| OsGRAS25 | LOC_Os05g40710 | UP | E(4.5) | 4.5 | UP | E(4.8) | 4.8 |
| OsGRAS26 | LOC_Os05g42130 | UP | IE(2.6) | 2.6 | DOWN | - | - |
| OsGRAS28 | LOC_Os06g01620 | DOWN | - | - | DOWN | - | - |
| OsCIGR1 | LOC_Os07g36170 | UP | IE(3.2) | 3.2 | DOWN | - | - |
| OsGRAS32 | LOC_Os07g38030 | DOWN | - | - | DOWN | - | - |
| OsCIGR2 | LOC_Os07g39470 | DOWN | - | - | DOWN | - | - |
| OsSHR1 | LOC_Os07g39820 | UP | IE(7.3) | 11.1 | DOWN | - | - |
| OsGRAS35 | LOC_Os07g40020 | UP | E(6.8) | 17.4 | DOWN | - | - |
| OsSCR1 | LOC_Os11g03110 | UP | E(3.8) | 5.7 | DOWN | - | - |
| ΨOsGRAS8 | LOC_Os11g04400 | DOWN |  |  | DOWN | - | - |
| OsGRAS39 | LOC_Os11g04570 | UP | IE(3.6) | 65.3 | DOWN | - | - |
| OsGRAS41 | LOC_Os11g06180 | UP | IE(2.4) | 4.1 | DOWN | - | - |
| ΨOsGRAS9 | LOC_Os11g11600 | UP | L(8.0) | 8 | DOWN | - | - |
| OsGRAS43 | LOC_Os11g31100 | UP | IE(3.5) | 3.5 | DOWN | - | - |
| OsGRAS44 | LOC_Os11g47870 | UP | IE(2.9) | 5.8 | DOWN | - | - |
| OsGRAS47 | LOC_Os11g47910 | - | - | - | DOWN | - | - |
| ΨOsGRAS10 | LOC_Os12g06540 | DOWN | - | - | DOWN | - | - |

| OsGRAS53 | LOC_Os12g38490 | DOWN | - | - | DOWN | - | - |
| --- | --- | --- | --- | --- | --- | --- | --- |
| Gene name | Locus number | Regulation<br>(up/down) | Type of<br>Response | Maximum<br>fold change | Regulation<br>(up/down) | Type of<br>Response | Maximum<br>fold change |
|  |  | **Root** |  |  | **Shoot** |  |  |
| **ABA** |  |  |  |  |  |  |  |
| OsGRAS1 | LOC_Os01g45860 | - | - | - | DOWN | - | - |
| OsGRAS2 | LOC_Os01g62460 | UP | IE(3.3) | 5.6 | DOWN | - | - |
| OsGRAS3 | LOC_Os01g65900 | UP | L(3.5) | 3.5 | DOWN | - | - |
| OsGRAS5 | LOC_Os01g71970 | UP | IE(6.2) | 12.2 | DOWN | - | - |
| OsGRAS7 | LOC_Os02g10360 | DOWN | - | - | DOWN | - | - |
| ΨOsGRAS2 | LOC_Os02g21685 | UP | IE(2.8) | 32.7 | UP | L(3.5) | 3.5 |
| OsGRAS8 | LOC_Os02g44360 | UP | IE(5.5) | 27.9 | DOWN | - | - |
| OsGRAS10 | LOC_Os02g45760 | UP | IE(2.4) | 16.7 | DOWN | - | - |
| OsGRAS11 | LOC_Os03g09280 | UP | L(4.1) | 4.1 | DOWN | - | - |
| OsGRAS12 | LOC_Os03g15680 | UP | IE(8.2) | 8.2 | UP | L(3.8) | 3.8 |
| OsSHR2 | LOC_Os03g31880 | UP | IE(4.5) | 14.3 | DOWN | - | - |
| ΨOsGRAS3 | LOC_Os03g37900 | UP | IE(2.8) | 2.8 | DOWN | - | - |
| ΨOsGRAS4 | LOC_Os03g40080 | UP | IE(2.1) | 2.1 | DOWN | - | - |
| OsGRAS15 | LOC_Os03g48450 | UP | IE(2.9) | 14.6 | DOWN | - | - |
| OsSLR1 | LOC_Os03g49990 | UP | IE(7.9) | 30.2 | DOWN | - | - |
| OsGRAS18 | LOC_Os03g51330 | UP | IE(2.8) | 3.4 | DOWN | - | - |
| OsGRAS19 | LOC_Os04g35250 | UP | L(3.5) | 11 | DOWN | - | - |
| ΨOsGRAS5 | LOC_Os04g37440 | UP | IE(2.9) | 2.9 | UP | IE(8.2) | 31.6 |
| OsGRAS20 | LOC_Os04g46860 | UP | IE(2.9) | 4.1 | DOWN | - | - |
| OsGRAS22 | LOC_Os04g50060 | UP | E(5.5) | 14.2 | DOWN | - | - |
| OsGRAS23 | LOC_Os05g31380 | DOWN | - | - | DOWN | - | - |
| OsGRAS24 | LOC_Os05g31420 | UP | IE(15.3) | 15.3 | UP | L(6.9) | 6.9 |
| OsGRAS25 | LOC_Os05g40710 | UP | IE(6.2) | 14.2 | UP | IE(3.0) | 10.5 |
| OsGRAS26 | LOC_Os05g42130 | UP | L(2.4) | 2.4 | DOWN | - | - |
| OsGRAS28 | LOC_Os06g01620 | DOWN | - | - | DOWN | - | - |
| OsCIGR1 | LOC_Os07g36170 | UP | IE(7.3) |  | DOWN | - | - |
| OsGRAS32 | LOC_Os07g38030 | UP | IE(2.09) |  | DOWN | - | - |
| OsCIGR2 | LOC_Os07g39470 | DOWN |  |  | DOWN | - | - |
| OsSHR1 | LOC_Os07g39820 | UP | IE(14.1) | 66.3 | DOWN | - | - |
| OsGRAS35 | LOC_Os07g40020 | UP | IE(15.9) | 15.9 | DOWN | - | - |
| OsSCR1 | LOC_Os11g03110 | UP | IE(6.7) | 34.3 | UP | L(2.4) | 2.4 |
| ΨOsGRAS8 | LOC_Os11g04400 | DOWN | - | - | DOWN | - | - |
| OsGRAS39 | LOC_Os11g04570 | UP | IE(12) | 101.4 | DOWN | - | - |
| OsGRAS41 | LOC_Os11g06180 | UP | IE(5) | 21.3 | DOWN | - | - |
| ΨOsGRAS9 | LOC_Os11g11600 | UP | L(2.7) | 2.7 | DOWN | - | - |
| OsGRAS43 | LOC_Os11g31100 | UP | IE(2.07) | 7.3 | DOWN | - | - |
| OsGRAS44 | LOC_Os11g47870 | UP | IE(19.2) | 19.2 | DOWN | - | - |
| OsGRAS47 | LOC_Os11g47910 |  | - | - | DOWN | - | - |
| ΨOsGRAS10 | LOC_Os12g06540 | DOWN | - | - | DOWN | - | - |
| OsGRAS53 | LOC_Os12g38490 | DOWN | - | - | DOWN | - | - |

Among the genes studied, some genes were observed to be expressed only under NaCl or ABA treatments at certain time points, whereas some were found to be expressive under both treatments. Such overlaps has been depicted in the form of Venn diagrams in Fig. S4. The corresponding list of genes clearly demonstrates that several genes were up/down-regulated simultaneously under both ABA and NaCl treatments at certain time points. In roots, the expression of 37.5% of the genes (IE type) overlapped under both stress treatments, while in shoots only *ΨOsGRAS5* (IE) was expressive.

### 3.7. Differential expression analysis of GRAS genes under biotic stress

We have studied the expression of the selected GRAS transcription factors in the leaf samples of rice infected with *Xoo* and *R. solani* pathogens that cause Bacterial Leaf Blight (BLB) and Sheath Blight (SB) diseases, respectively (Fig. 6). Six genes were upregulated in BLB of which five (*OsGRAS1, OsGRAS18, OsCIGR2, ΨOsGRAS9, OsGRAS53*) showed low expression while one gene (*OsCIGR1*) was highly upregulated upto 57 folds. More genes were upregulated in SB infected leaves compared to the BLB treated ones. Out of the thirty expressed genes in SB infected leaves, only twelve showed very high expression levels while the rest of the genes exhibited low to moderate expression. *OsGRAS2, ΨOsGRAS3, OsGRAS19, OsGRAS20, OsGRAS23* and *OsSHR1* were expressed by ≥100 folds under the SB treatment. A total of 22 genes in BLB and three in SB treated samples were downregulated. Those that were downregulated in SB treated samples (*OsSHR2*, *OsGRAS24* and *OsGRAS43*) were also downregulated in BLB treated leaves. Twelve genes under BLB and seven under SB showed no changes in their expression levels.

**Figure 6:**
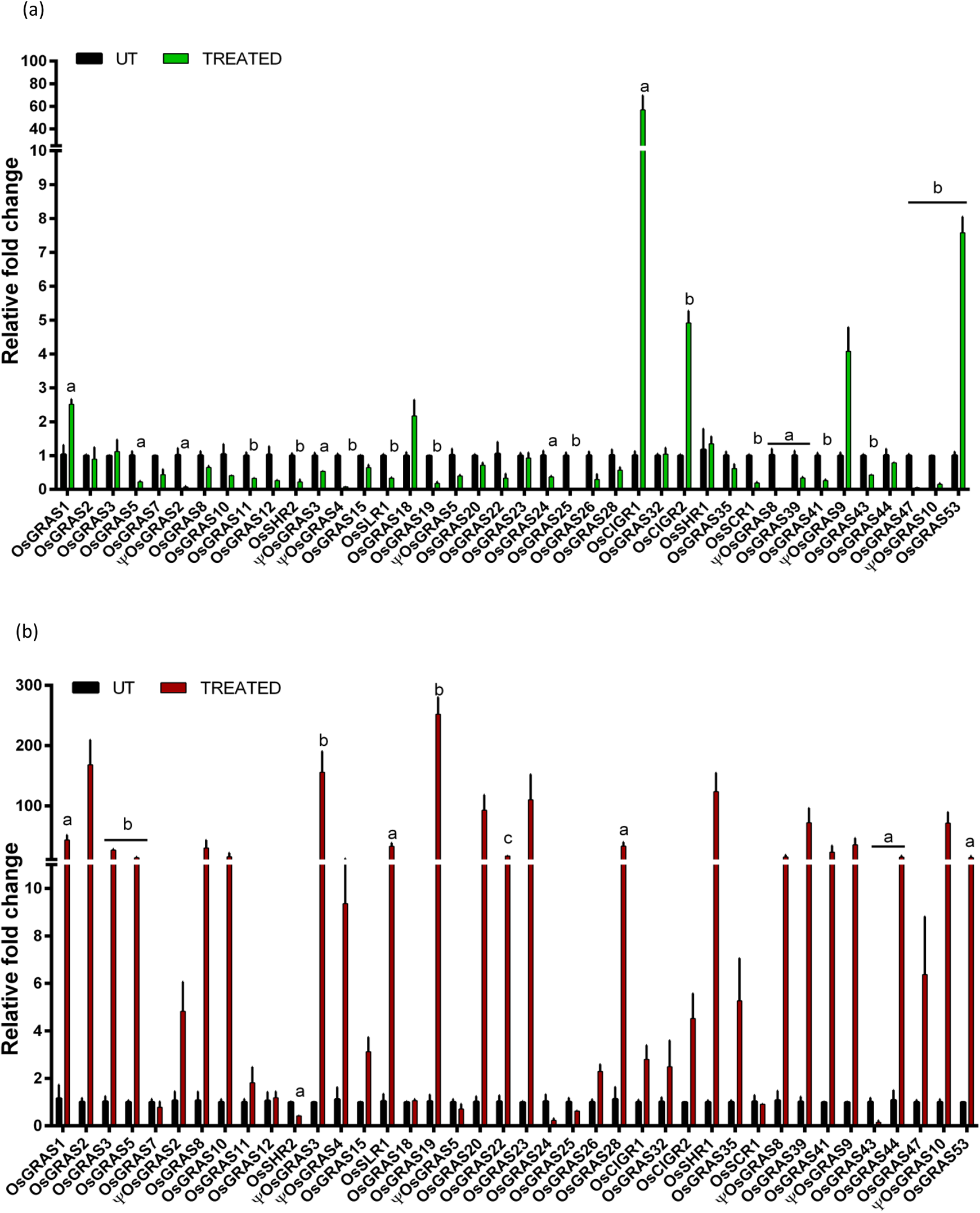
Quantitative real-time expression analysis of GRAS genes under biotic stress. Expression analysis of GRAS genes under the infection of *Xanthomonas oryzae* pv. *oryzae* causing bacterial leaf blight (6a) and *Rhizoctonia solani* causing sheath blight (6b) were studied. The genes were double normalized using rice *actin* and *tubulin* as internal reference genes and the C_T_ values untreated samples by ΔΔC_T_ method. One way ANOVA was performed on the data and a represents *P*<0.05, b represents *P*<0.025 and c represents *P*<0.001

### 3.8. Native expression analysis of GRAS genes in various tissues at specific developmental stages in rice

In order to study the native expression patterns of GRAS transcription factors in different tissues of the rice plant, we performed qRT-PCR analysis of 13 different tissues, which included shoot, root, root-shoot transition, flag leaves, flower, spikes and grain of mature 20 d old plants (after shifting to greenhouse), shoot and root of 7 d old seedlings, 3 d old plumule and radicle, embryo and endosperm of 16 h germinating seeds. The mean values were used to plot a heat map (Fig. 7).

**Figure 7:**
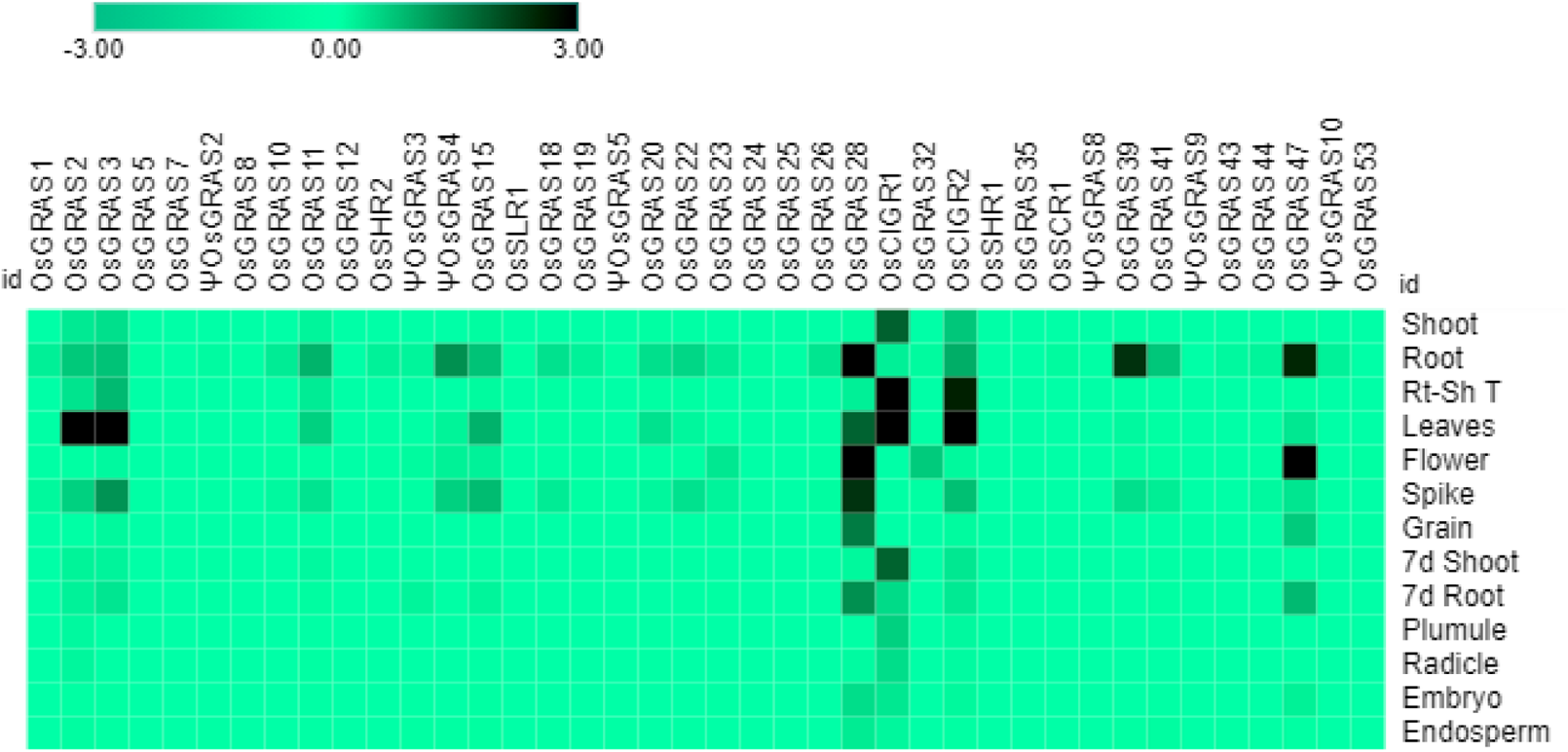
Native expression analysis of GRAS genes. Heat map representing the spatial expression pattern of GRAS genes under 13 different developmental stages of rice. The map was generated using MORPHEUS program. The data was single normalized using rice *actin* as the internal reference gene.

Expression analysis showed a conspicuous downregulation of all genes in most of the tissues particularly in plumule, radicle, embryo and the endosperm. The number of downregulated genes in other tissues are: 39 genes in 7 d shoot, 38 genes in 20 d shoot, grain and 7d root, and 37genes in root-shoot transition and flower followed by 36 genes in spikes, 34 genes in leaves and 30 genes in 20 d root. Out of 40 selected genes, only seven were expressed in certain tissues. *OsGRAS2* and *OsGRAS3* were upregulated only in mature leaves, *OsGRAS28* in 20 d root, flower and spike, *OsCIGR1* in root-shoot transition and leaves, *OsGRAS39* in 20 d root and *OsGRAS47* in 20 d root and flower. Out of these seven genes, five were upregulated either in the roots or in the root-shoot transition region indicating the preference of GRAS genes towards expression in the root tissue. This is also in accordance with the expression analysis under abiotic stress condition, where the genes were highly expressive in roots rather than in the shoot tissue. Three out of seven mildly expressive genes were upregulated in flower and spike of 20 d old plants with none of them expressing in the grain. *OsGRAS39*, which was upregulated in root tissues under native conditions, is highly expressive in roots under abiotic stress conditions also responding immediately after the application of stress treatment. This might be an indication of its tissue specificity and its potential as a stress tolerance transcription factor gene.

## 4. Discussion

Being sessile, plants cannot escape the onslaught from environmental stresses like cold, heat, drought etc., nor can they avoid harmful interactions with microorganisms like fungi and bacteria (Lin et al., 2017). Such adversities impose a threat to agricultural productivity and sustainability. In order to support the burgeoning global population, the development of stress tolerant crops is of utmost importance (Cushman and Bohnert, 2000). Characterization of insertional mutants is an important functional genomics based method of identifying novel genes responsible for inducing stress tolerance in crop plants (Cushman and Bohnert, 2000). TF genes are of particular importance in this context as they act upstream in the pathway(s) and control the expression of several genes working under their control. Because of this, the manipulation undertaken using TF genes as ‘Master’ genes would render the plant more accommodative towards the particular stress under consideration. Our previous studies have identified several key players for stress tolerance in *indica* rice variety via enhancer based activation tagging method. A GRAS transcription factor gene, *ΨOsGRAS4* was one of the important genes that was identified in the study along with others for enhanced water use efficiency associated with enhanced photosynthetic efficiency (Moin et al., 2016a).

### 4.1. Evolutionary relationships, gene organization and protein properties of rice GRAS genes

Plants use certain acclimation and adaptive measures to cope up with the impending stress, which is mostly modulated through the action of hormones and regulators (Lin et al., 2017). Thus understanding the expression patterns of GRAS family of genes, which play a key role in gibberellin signaling and their spatio-temporal regulation help us identify candidate genes for improving the endogenous defence ability of plants, particularly rice in the present context. In this study, we have shortlisted important GRAS genes responsible for abiotic and biotic stress tolerance. We have also studied the *in-silico* properties of these genes and have correlated them with our expression data.

According to the published evidence that is available (Liu and Widmer, 2014), 60 GRAS genes were reported in the rice genome, which are distributed on all the twelve chromosomes except chromosome numbers 8 and 9. Highest number of gene density was observed on chromosome 11. We have selected 40 genes for our study, drawing one member representing each paralogous group. The availability of high quality genomic sequences enabled us to get an insight into the phylogenetic, genomic and protein properties of the GRAS genes. In our analysis, we have classified the genes into 14 subfamilies (Cenci and Rouard, 2017) of which LISCL constituted the maximum number of genes. However, most expressive genes belonged to SCL3, SHR1, DELLA, HAM and PAT subfamilies. The ten MEME-identified motifs were categorized into five conserved C-terminal GRAS motifs. Genes belonging to the same subfamily exhibited similar motif arrangements, but this varied within the subfamilies, which might be due to the diverse biological functions of GRAS genes. This group of proteins were reported to have originated in bacteria, which later expanded into eukaryotic genomes via horizontal gene transfer and repeated duplication events with the possible retroposition of intronless genes (Huang et al., 2015). Our genomic organization study revealed 31 *OsGRAS* genes out of 40 to be intronless and this observation is in line with previous studies. Several interacting metallic and non-metallic ligands associated with GRAS genes along with their hydrophilic nature (as indicated by the GRAVY index) indicated their involvement in cell signaling, catalysis and protein-protein interactions (Ulucan et al., 2014; Jing et al., 2017). The majority of the genes were observed to have a *p*I less than seven and were found to be rich in negatively charged amino acids like glutamic acid and aspartic acid. This makes the interactions of GRAS proteins very specific as proteins with low *p*I tend to minimize the chances of non-specific interactions with nucleic acids and other acidic proteins (Takakura et al., 2016). All GRAS genes have at least one GRAS domain, but some were found to have two or possess a DELLA domain, which is known to have important role in gibberellic acid signaling (Urbanova and Metzger, 2018).

### 4.2. Differential expression patterns of *OsGRAS* genes and their spatio-temporal regulation

Based on external cues, spatio-temporal regulation of gene transcription is required to control the concentration of particular transcripts and proteins in the cells for their adjustment to the environmental changes. Most of the GRAS genes were observed to be upregulated in roots with 55-60% of them showing IE type of gene expression. In plants, stress responses can be divided broadly into early and late response types. Early responsive genes are expressed within minutes of stress induction and this provides protection and repair from the initial stress. Such response “alarms” the plant to prepare for further stress tolerance or avoidance. On the other hand, late responsive genes are mostly involved in protein synthesis that regulates downstream genes, thereby responding to the “adaptation” part of stress mediation (Bahrami and Drablos, 2016; Lin et al., 2017). *ΨOsGRAS9* was observed to be late expressive under both treatments in root, indicating that it might have an important role in subsequent steps of stress amelioration. Interestingly, 50-60% of IE expressed genes continued to express till the end point of the treatment. Among them, *OsGRAS39* was the only gene that was observed to be highly expressive under both NaCl and ABA treatments in roots. Apart from this, *ΨOsGRAS2, OsSHR1* and *OsSCR1* continued their expression till 60 h under ABA treatment. This probably indicates that these constitute an important set of genes required by the plant throughout for stress remediation. Others like *ΨOsGRAS3, ΨOsGRAS4, OsCIGR1* and *OsGRAS32* under ABA and *OsCIGR1* under NaCl were the genes induced initially (IE type), which either stopped expressing or got downregulated at subsequent time points. These genes are probably required for initial stress responses whose function is later on taken up by other downstream genes in the signaling cascade. *ΨOsGRAS5* (IE type) was found to be the only moderately expressive gene in shoot under both stress conditions. Since root is the first organ to perceive the stress signal, it induces a signaling cascade that extends towards shoot. Such preferential expression of *GRAS* genes in roots over shoots indicates their important role in stress tolerance (Janiak et al., 2016). Also, previous studies indicated that probably the role of these genes in pattern formation and signal transduction enables them to be more expressive in roots (Pysh et al., 1999).

Rice productivity is severely hampered by BLB and SB diseases. BLB infection during tillering stage can cause a yield decline of 20-40%, while it can reduce crop productivity by 50% at a younger stage. Upto 45% yield losses in rice are caused by SB infections (Chukwu et al., 2019; Singh et al., 2019). Thus, identification of key genes and understanding their expression patterns are important for developing tolerant varieties of rice for these diseases. Under BLB treatment, only six genes (15%) were expressive compared to thirty (75%) expressive genes under SB infection. Quite a number of genes were expressive under SB infection and both abiotic stress conditions. Noteworthy among them are *OsGRAS39*, *OsGRAS8*, *OsSHR1* and *OsSLR1*. Thus, these genes can be considered as to be quite important as they are expressive under both stress conditions with important roles in disease resistance. Majority of the genes were highly expressive in ABA treated roots and SB infection. The presence of multiple stress responsive elements in their putative promoter regions are corroborated by our expression data and these observations indicate their probable roles in improving plant defence against biotic and abiotic stress.

Proteins belonging to DELLA subfamily of GRAS transcription factors are known to be negative regulators of seed germination as bioactive gibberellic acid causes proteasomal degradation of such proteins for gibberellin signaling to occur (Urbanova and Metzger, 2018). This explains the downregulation of all genes in plumule, radicle, embryo and endosperm. *OsGRAS39* was expressive in roots under native as well as abiotic stress conditions indicating its tissue specificity. This gene belongs to SCL3 subfamily, which is known for modulating GA signaling in roots via protein-protein interactions (Weng et al., 2020). Hence, high expression of *OsGRAS39* in roots under all stress conditions can be further exploited for its potential role in stress tolerance.

GRAS gene family has been studied extensively in many plant species, but we have successfully provided a backdrop based on which future exploration on rice GRAS genes can be done. The differential expression patterns of these genes indicates their importance in stress remediation. Our study provides an insight into the role of GRAS genes in stress tolerance along with their spatio-temporal regulation. Based on this report, it would be possible to pick up important genes that can be further manipulated to develop stress tolerant varieties of rice and other related crops.

## Author contribution statement

PBK and MD designed the experiments and prepared the manuscript. MD performed the experiments. AS helped in the qRT-PCR experiments and analysis. MM identified GRAS gene from activation tagged lines and conceived the initial idea. PBK supervised the work.

## Acknowledgements

MD is grateful to UoH BBL and UGC for research fellowship and contingency and to DBT for funding the rice activation tagging project. She is also grateful to Prof. S. Dayananda, Dean School of Life Sciences, UoH for his help. MD and AS are grateful to DBT for providing research fellowship and contingency. MD is grateful to Dr. M.S. Madhav, Department of Biotechnology, Indian Institute of Rice Research, Hyderabad, India, for providing the infected rice samples and wild type BPT-5204 seeds. PBK acknowledges the National Academy of Sciences-India for the award of NASI-Platinum Jubilee Senior Scientist position.

## Conflict of interest

The authors hereby declare that the research has been performed without any financial and commercial conflict of interest.

## Notes

### Competing Interest Statement

The authors have declared no competing interest.

## References

Ambawat, Supriya, Poonam Sharma, Neelam R. Yadav, and Ram C. Yadav. 2013. “MYB Transcription Factor Genes as Regulators for Plant Responses: An Overview.” Physiology and Molecular Biology of Plants 19(3):307–21.

Bahrami, Shahram, and Finn Drabløs. 2016. “Gene Regulation in the Immediate-Early Response Process.” Advances in Biological Regulation 62:37–49.

Bernard, Virginie, Véronique Brunaud, and Alain Lecharny. 2010. “TC-Motifs at the TATA-Box Expected Position in Plant Genes: A Novel Class of Motifs Involved in the Transcription Regulation.” BMC Genomics 11(1):1–15.

Bolle, Cordelia. 2004. “The Role of GRAS Proteins in Plant Signal Transduction and Development.” Planta 218(5):683–92.

Bolle, Cordelia, Csaba Koncz, and Nam Hai Chua. 2000. “PAT1, a New Member of the GRAS Family, Is Involved in Phytochrome A Signal Transduction.” Genes and Development 14(10):1269–78.

Cenci, Alberto, and Mathieu Rouard. 2017. “Evolutionary Analyses of GRAS Transcription Factors in Angiosperms.” Frontiers in Plant Science 8(March):1–15.

Choi, Hyung In, Jung Hee Hong, Jin Ok Ha, Jung Youn Kang, and Soo Young Kim. 2000. “ABFs, a Family of ABA-Responsive Element Binding Factors.” Journal of Biological Chemistry 275(3):1723–30.

Chukwu, S. C., · M Y Rafii, · S I Ramlee, · S I Ismail, · M M Hasan, Y. A. Oladosu, · U G Magaji, Ibrahim Akos, and · K K Olalekan. 2019. “Bacterial Leaf Blight Resistance in Rice: A Review of Conventional Breeding to Molecular Approach.” Molecular Biology Reports 46:1519–32.

Cushman, John C., and Hans J. Bohnert. 2000. “Genomic Approaches to Plant Stress Tolerance.” Current Opinion in Plant Biology 3(2):117–24.

Dhatterwal, Pinky, Samyadeep Basu, Sandhya Mehrotra, and Rajesh Mehrotra. 2019. “Genome Wide Analysis of W-Box Element in Arabidopsis Thaliana Reveals TGAC Motif with Genes down Regulated by Heat and Salinity.” Scientific Reports 9(1):1–8.

Ding, Shuangcheng, Fengyu He, Wenlin Tang, Hewei Du, and Hongwei Wang. 2019. “Identification of Maize Cc-Type Glutaredoxins That Are Associated with Response to Drought Stress.” Genes 10(8).

Dutta, Mouboni, Mazahar Moin, Anusree Saha, Dibyendu Dutta, Achala Bakshi, and P. B. Kirti. 2021. “Gain-of-Function Mutagenesis through Activation Tagging Identifies XPB2 and SEN1 Helicase Genes as Potential Targets for Drought Stress Tolerance in Rice.” Theoretical and Applied Genetics.

Guo, Yuyu, Hongyu Wu, Xiang Li, Qi Li, Xinyan Zhao, Xueqing Duan, Yanrong An, Wei Lv, and Hailong An. 2017. “Identification and Expression of GRAS Family Genes in Maize (Zea Mays L.)” edited by M. Sun. PLOS ONE 12(9):e0185418.

Helariutta, Yrjo, Hidehiro Fukaki, Joanna Wysocka-Diller, Keiji Nakajima, Jee Jung, Giovanni Sena, Marie Theres Hauser, and Philip N. Benfey. 2000. “The SHORT-ROOT Gene Controls Radial Patterning of the Arabidopsis Root through Radial Signaling.” Cell 101(5):555–67.

Huang, Wei, Zhiqiang Xian, Xia Kang, Ning Tang, and Zhengguo Li. 2015. “Genome-Wide Identification, Phylogeny and Expression Analysis of GRAS Gene Family in Tomato.” BMC Plant Biology 15(1):1–18.

Hwang, Jung Eun, Joon Ki Hong, Chan Ju Lim, Huan Chen, Jihyun Je, Kyung Ae Yang, Dool Yi Kim, Young Ju Choi, Sang Yeol Lee, and Chae Oh Lim. 2010. “Distinct Expression Patterns of Two Arabidopsis Phytocystatin Genes, AtCYS1 and AtCYS2, during Development and Abiotic Stresses.” Plant Cell Reports 29(8):905–15.

Janiak, Agnieszka, Mirosław Kwasniewski, and Iwona Szarejko. 2016. “Gene Expression Regulation in Roots under Drought.” Journal of Experimental Botany 67(4):1003–14.

Jing, Zhifeng, Rui Qi, Chengwen Liu, and Pengyu Ren. 2017. “Study of Interactions between Metal Ions and Protein Model Compounds by Energy Decomposition Analyses and the AMOEBA Force Field.” Journal of Chemical Physics 147(16):161733.

Kelley, Lawrence A., Stefans Mezulis, Christopher M. Yates, Mark N. Wass, and Michael J. E. Sternberg. 2015. “The Phyre2 Web Portal for Protein Modeling, Prediction and Analysis.” Nature Protocols 10(6):845–58.

Lin, Chung-Wen, Li-Yao Huang, Chao-Li Huang, Yong-Chuan Wang, Pei-Hsuan Lai, Hao-Ven Wang, Wen-Chi Chang, Tzen-Yuh Chiang, and Hao-Jen Huang. 2017. “Common Stress Transcriptome Analysis Reveals Functional and Genomic Architecture Differences Between Early and Delayed Response Genes.” Plant and Cell Physiology 58(3):pcx002.

Liu, Limin, Xiaomei Zhang, Fulu Chen, Asia Adam Elzamzami Mahi, Xiaoxia Wu, Qingshan Chen, and Yong Fu Fu. 2017. “Analysis of Promoter Activity Reveals That GmFTL2 Expression Differs from That of the Known Flowering Locus T Genes in Soybean.” Crop Journal 5(5):438–48.

Liu, Xuanyu, and Alex Widmer. 2014. “Genome-Wide Comparative Analysis of the GRAS Gene Family in Populus, Arabidopsis and Rice.” Plant Molecular Biology Reporter 32(6):1129–45.

Livak, Kenneth J., and Thomas D. Schmittgen. 2001. “Analysis of Relative Gene Expression Data Using Real-Time Quantitative PCR and the 2-ΔΔCT Method.” Methods.

Ma, Hong-Shuang, Dan Liang, Peng Shuai, Xin-Li Xia, and Wei-Lun Yin. 2010. “The Salt- and Drought-Inducible Poplar GRAS Protein SCL7 Confers Salt and Drought Tolerance in Arabidopsis Thaliana.” Journal of Experimental Botany 61(14):4011–19.

Moin, Mazahar, Achala Bakshi, Anusree Saha, Mouboni Dutta, Sheshu M. Madhav, and P. B. Kirti. 2016. “Rice Ribosomal Protein Large Subunit Genes and Their Spatio-Temporal and Stress Regulation.” Frontiers in Plant Science 7(AUG2016):1–20.

Moin, Mazahar, Achala Bakshi, Anusree Saha, M. Udaya Kumar, Attipalli R. Reddy, K. V. Rao, E. A. Siddiq, and P. B. Kirti. 2016. “Activation Tagging in Indica Rice Identifies Ribosomal Proteins as Potential Targets for Manipulation of Water-Use Efficiency and Abiotic Stress Tolerance in Plants.” Plant Cell and Environment 39(11):2440–59.

Narusaka, Yoshihiro, Kazuo Nakashima, Zabta K. Shinwari, Yoh Sakuma, Takashi Furihata, Hiroshi Abe, Mari Narusaka, Kazuo Shinozaki, and Kazuko Yamaguchi-Shinozaki. 2003. “Interaction between Two Cis-Acting Elements, ABRE and DRE, in ABA-Dependent Expression of Arabidopsis Rd29A Gene in Response to Dehydration and High-Salinity Stresses.” Plant Journal 34(2):137–48.

Oñate-Sánchez, Luis, and Karam B. Singh. 2002. “Identification of Arabidopsis Ethylene-Responsive Element Binding Factors with Distinct Induction Kinetics after Pathogen Infection.” Plant Physiology 128(4):1313–22.

Pysh, Leonard D., Joanna W. Wysocka-Diller, Christine Camilleri, David Bouchez, and Philip N. Benfey. 1999. “The GRAS Gene Family in Arabidopsis: Sequence Characterization and Basic Expression Analysis of the SCARECROW-LIKE Genes.” Plant Journal 18(1):111–19.

Saha, Anusree, Shubhajit Das, Mazahar Moin, Mouboni Dutta, Achala Bakshi, M. S. Madhav, and P. B. Kirti. 2017. “Genome-Wide Identification and Comprehensive Expression Profiling of Ribosomal Protein Small Subunit (RPS) Genes and Their Comparative Analysis with the Large Subunit (RPL) Genes in Rice.” Frontiers in Plant Science 8(September):1–21.

Sakai, Tatsuya, Yohsuke Takahashi, and Toshiyuki Nagata. 1996. “Analysis of the Promoter of the Auxin-Inducible Gene, ParC, of Tobacco.” Plant and Cell Physiology 37(7):906– 13.

Shen, Zijie, Yuan Lin, and Quan Zou. 2020. “Transcription Factors–DNA Interactions in Rice: Identification and Verification.” Briefings in Bioinformatics 21(3):946–56.

Sidhu, Navjot Singh, Gomsie Pruthi, Sahildeep Singh, Ritika Bishnoi, and Deepak Singla. 2020. “Genome-Wide Identification and Analysis of GRAS Transcription Factors in the Bottle Gourd Genome.” Scientific Reports 10(1):14338.

Singh, Pooja, Purabi Mazumdar, Jennifer Ann Harikrishna, and Subramanian Babu. 2019. “Sheath Blight of Rice: A Review and Identification of Priorities for Future Research.” Planta 250(5):1387–1407.

Stuurman, Jeroen, Fabienne Jäggi, and Cris Kuhlemeier. 2002. “Shoot Meristem Maintenance Is Controlled by a GRAS-Gene Mediated Signal from Differentiating Cells.” Genes and Development 16(17):2213–18.

Sun, Xiaolin, William T. Jones, and Erik H. A. Rikkerink. 2012. “GRAS Proteins: The Versatile Roles of Intrinsically Disordered Proteins in Plant Signalling.” Biochemical Journal 442(1):1–12.

Takakura, Yoshimitsu, Kozue Sofuku, Masako Tsunashima, and Shigeru Kuwata. 2015. “Lentiavidins: Novel Avidin-like Proteins with Low Isoelectric Points from Shiitake Mushroom (Lentinula Edodes).”

Tian, Chaoguang, Ping Wan, Shouhong Sun, Jiayang Li, and Mingsheng Chen. 2004. *Genome-Wide Analysis of the GRAS Gene Family in Rice and Arabidopsis*.

To, Vinh Trieu, Qi Shi, Yueya Zhang, Jin Shi, Chaoqun Shen, Dabing Zhang, and Wenguo Cai. 2020. “Genome-Wide Analysis of the Gras Gene Family in Barley (Hordeum Vulgare L.).” Genes 11(5):553.

Tran, Lam Son Phan, Kazuo Nakashima, Yoh Sakuma, Sean D. Simpson, Yasunari Fujita, Kyonoshin Maruyama, Miki Fujita, Motoaki Seki, Kazuo Shinozaki, and Kazuko Yamaguchi-Shinozaki. 2004. “Isolation and Functional Analysis of Arabidopsis Stress-Inducible NAC Transcription Factors That Bind to a Drought-Responsive Cis-Element in the Early Responsive to Dehydration Stress 1 Promoter.” Plant Cell 16(9):2481–98.

Ulucan, Ozlem, Tanushree Jaitly, and Volkhard Helms. 2014. “Energetics of Hydrophilic Protein-Protein Association and the Role of Water.” Journal of Chemical Theory and Computation 10(8):3512–24.

Urbanova, Terezie, and Gerhard Leubner-Metzger. 2018. “Gibberellins and Seed Germination.” Pp. 253–84 in Annual Plant Reviews online. Chichester, UK: John Wiley & Sons, Ltd.

Wang, Yin, Guan Jun Liu, Xiu Feng Yan, Zhi Gang Wei, and Zhi Ru Xu. 2011. “MeJA-Inducible Expression of the Heterologous JAZ2 Promoter from Arabidopsis in Populus Trichocarpa Protoplasts.” Journal of Plant Diseases and Protection 118(2):69–74.

Wass, Mark N., Lawrence A. Kelley, and Michael J. E. Sternberg. 2010. “3DLigandSite: Predicting Ligand-Binding Sites Using Similar Structures.” Nucleic Acids Research 38(SUPPL. 2):W469–73.

Wei, Qiang, Huiming Cao, Zhongru Li, Benke Kuai, and Yulong Ding. 2013. “Identification of an AtCRN1-like Chloroplast Protein BeCRN1 and Its Distinctive Role in Chlorophyll Breakdown during Leaf Senescence in Bamboo (Bambusa Emeiensis ‘Viridiflavus’).” Plant Cell, Tissue and Organ Culture 114(1):1–10.

Weng, Chun Yue, Mo Han Zhu, Zhi Qiang Liu, and Yu Guo Zheng. 2020. “Integrated Bioinformatics Analyses Identified SCL3-Induced Regulatory Network in Arabidopsis Thaliana Roots.” Biotechnology Letters 42(6):1019–33.

Whitbred, J. M., and M. A. Schuler. 2000. “Molecular Characterization of CYP73A9 and CYP82A1 P450 Genes Involved in Plant Defense in Pea.” Plant Physiology 124(1):47– 58.

Xu, Kai, Shoujun Chen, Tianfei Li, Xiaosong Ma, Xiaohua Liang, Xuefeng Ding, Hongyan Liu, and Lijun Luo. 2015. “OsGRAS23, a Rice GRAS Transcription Factor Gene, Is Involved in Drought Stress Response through Regulating Expression of Stress-Responsive Genes.” BMC Plant Biology 15(1):1–13.

Xu, Wei, Zexi Chen, Naeem Ahmed, Bing Han, Qinghua Cui, and Aizhong Liu. 2016. “Genome-Wide Identification, Evolutionary Analysis, and Stress Responses of the GRAS Gene Family in Castor Beans.” International Journal of Molecular Sciences 17(7).

Xu, Weirong, Yihe Yu, Qi Zhou, Jiahua Ding, Lingmin Dai, Xiaoqing Xie, Yan Xu, Chaohong Zhang, and Yuejin Wang. 2011. “Expression Pattern, Genomic Structure, and Promoter Analysis of the Gene Encoding Stilbene Synthase from Chinese Wild Vitis Pseudoreticulata.” Journal of Experimental Botany 62(8):2745–61.

Yin, Xiangjing, Li Huang, Xiuming Zhang, Chunlei Guo, Hao Wang, Zhi Li, and Xiping Wang. 2017. “Expression Patterns and Promoter Characteristics of the Vitis Quinquangularis VqSTS36 Gene Involved in Abiotic and Biotic Stress Response.” Protoplasma 254(6):2247–61.

Zeng, Xu, Hong Ling, Xiaomei Chen, and Shunxing Guo. 2019. “Genome-Wide Identification, Phylogeny and Function Analysis of GRAS Gene Family in Dendrobium Catenatum (Orchidaceae).” Gene 705(151):5–15.

Zhang, Bin, J. Liu, Zhao E. Yang, Er Y. Chen, Chao J. Zhang, Xue Y. Zhang, and Fu G. Li. 2018. “Genome-Wide Analysis of GRAS Transcription Factor Gene Family in Gossypium Hirsutum L.” BMC Genomics 19(1):1–12.

Zhang, Dapeng, Lakshminarayan M. Iyer, and L. Aravind. 2012. “Bacterial GRAS Domain Proteins Throw New Light on Gibberellic Acid Response Mechanisms.” Bioinformatics 28(19):2407–11.

Zhang, Hailing, Yingping Cao, Chen Shang, Jikai Li, Jianli Wang, Zhenying Wu, Lichao Ma, Tianxiong Qi, Chunxiang Fu, Zetao Bai, and Baozhong Hu. 2017. “Genome-Wide Characterization of GRAS Family Genes in Medicago Truncatula Reveals Their Evolutionary Dynamics and Functional Diversification” edited by H. Luo. PLOS ONE 12(9):e0185439.

Zhang, Huiling, Xijuan Zhao, Juping Zhang, Bo Yang, Yihe Yu, Tengfei Liu, Bihua Nie, and Botao Song. 2020. “Functional Analysis of an Anthocyanin Synthase Gene StANS in Potato.” Scientia Horticulturae 272:109569.

