## Supplementary material for "Genome-wide identification, expression and bioinformatic analyses of GRAS transcription factor genes in rice"

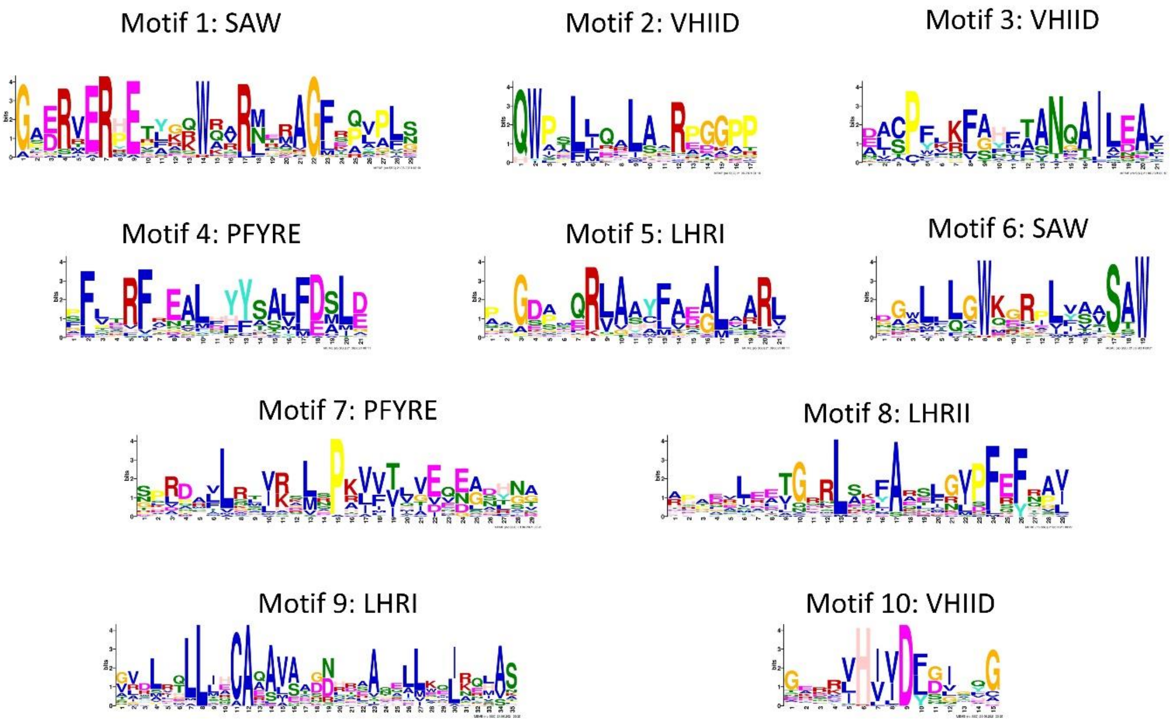


Figure S1: Logos representing each MEME identified motif. A total of 10 motifs were identified, which were classified according to the conserved motifs of GRAS domain. Each logo represents the conserved sequence of that motif. The height of each amino acid in a logo indicates its frequency at that position.

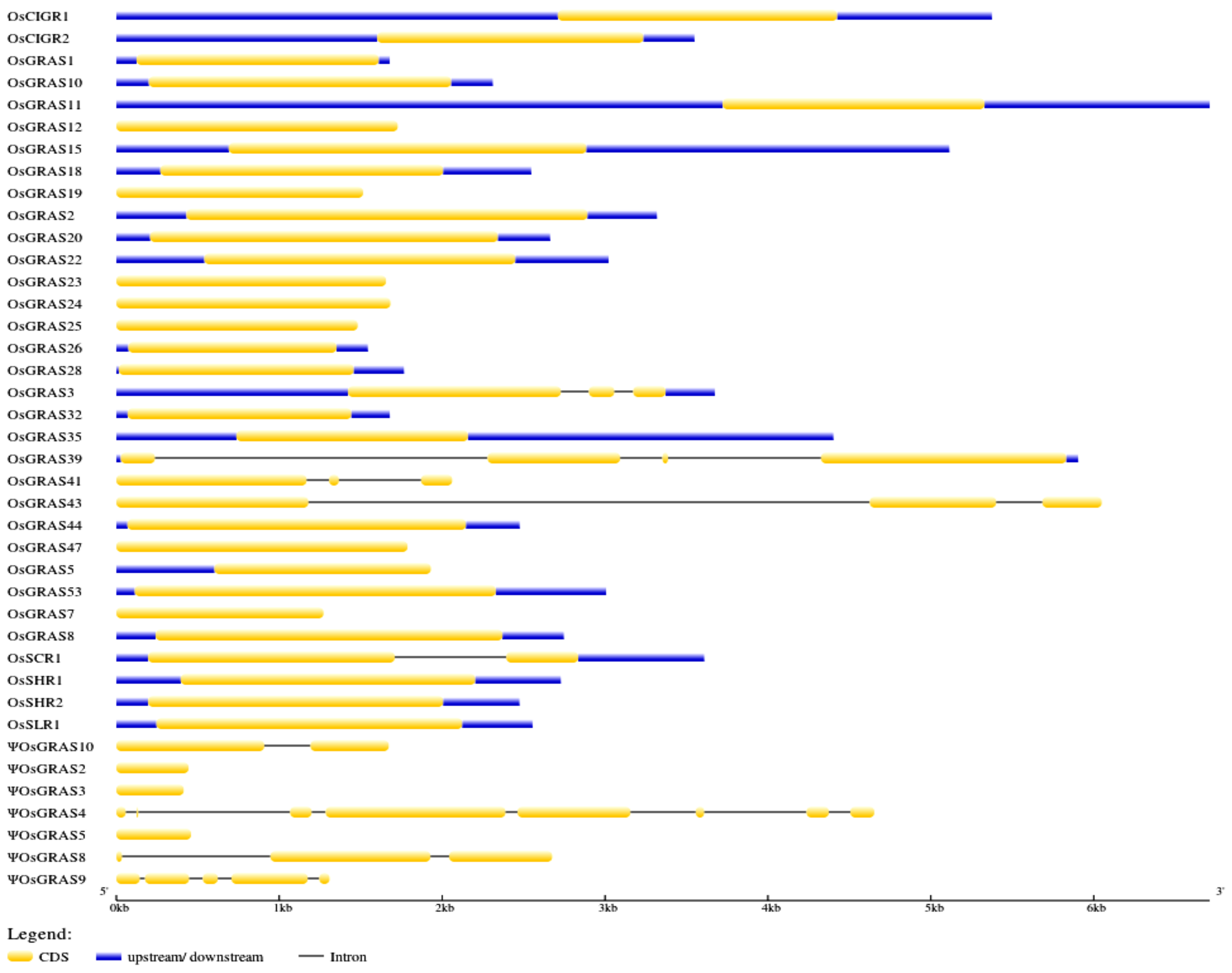


Figure S2: Figure representing the genetic organisation of rice GRAS genes under study (developed through Gene Structure Display Server (GSDSv2)). The yellow coded region indicates the coding sequence and the blue region indicates the untranslated regions. The black line corresponds to the intronic sequences.

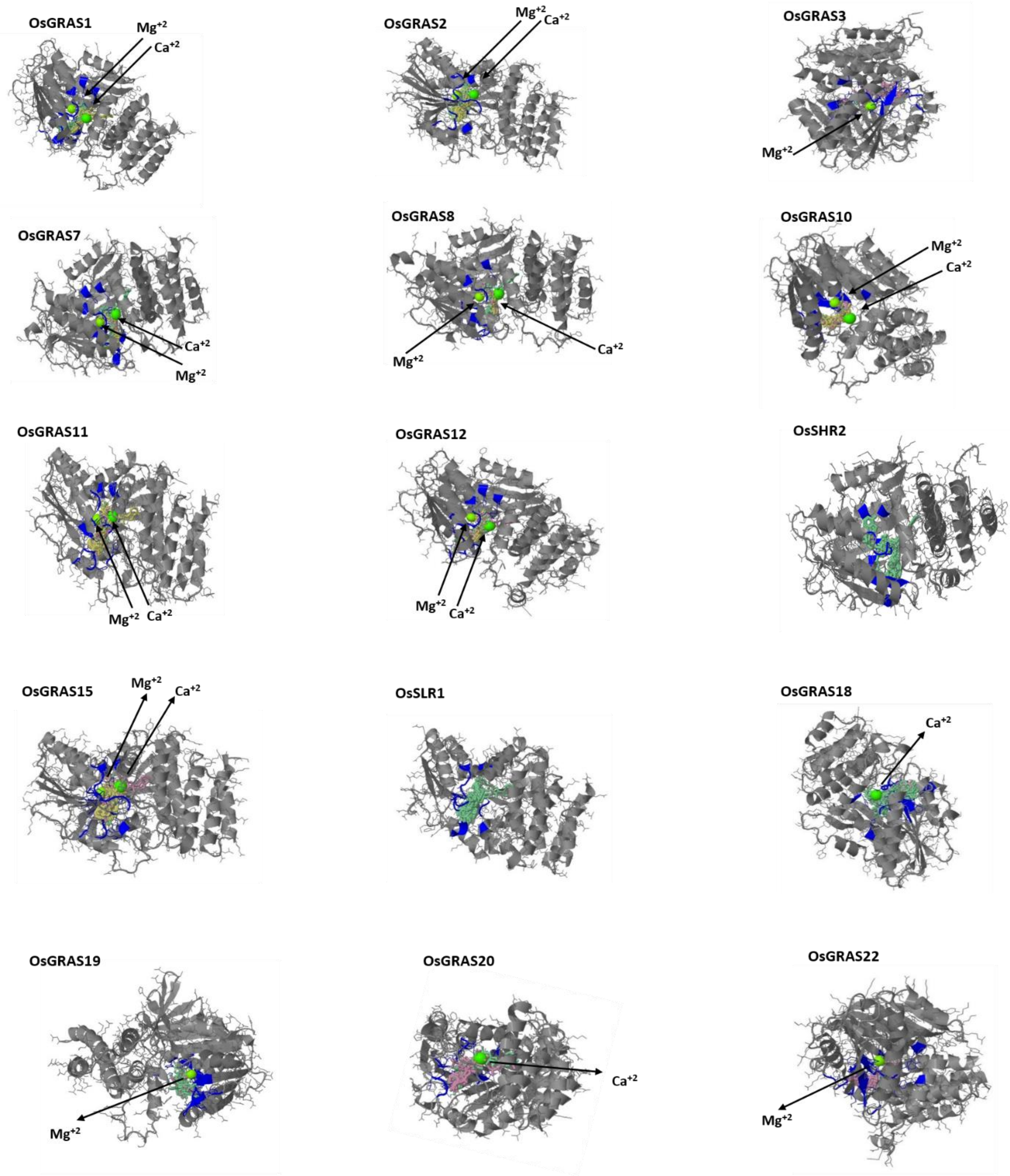

OsGRAS1
Mg+2
Ca+2
OsGRAS2
Mg+2
Ca+2
OsGRAS3
Mg+2
OsGRAS7
Ca+2
Mg+2
OsGRAS8
Mg+2
Ca+2
OsGRAS10
Mg+2
Ca+2
OsGRAS11
Mg+2
Ca+2
OsGRAS12
Mg+2
Ca+2
OsSHR2
OsGRAS15
Mg+2
Ca+2
OsSLR1
OsGRAS18
Ca+2
OsGRAS19
Mg+2
OsGRAS20
Ca+2
OsGRAS22
Mg+2

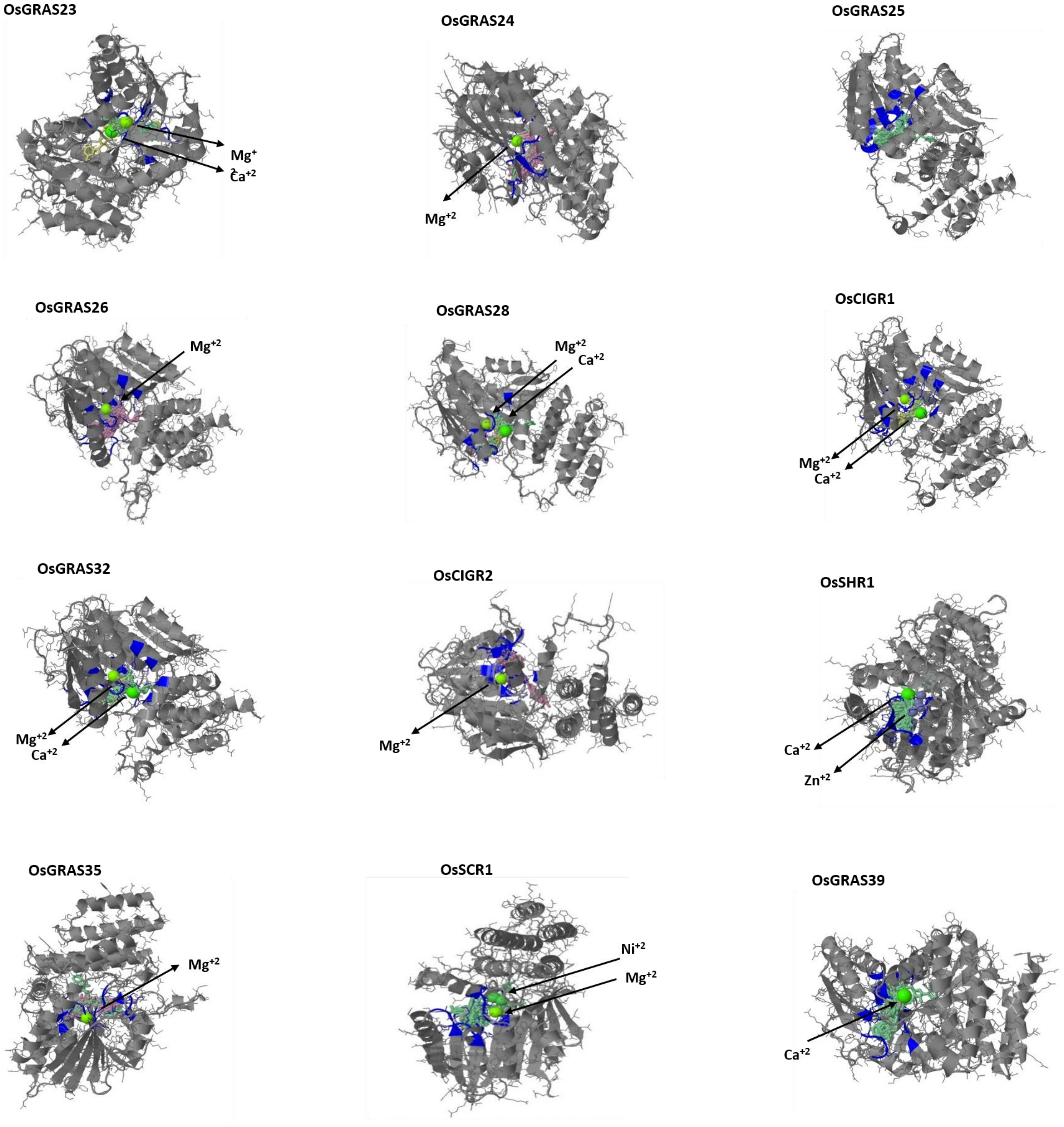


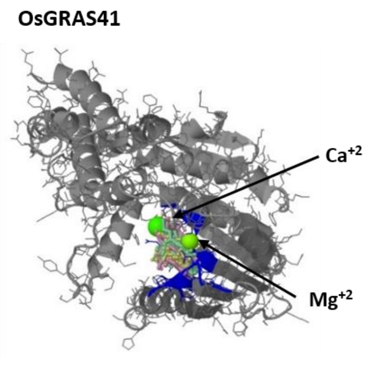


Figure S3: Three dimensional structures of 28 GRAS proteins along with their interacting ligands as predicted by 3Dligand site and Phyre2 program. The blue labelled region indicates the point of interaction with the metallic and non-metallic ligands. The metallic ligands are labelled. These include $Ca^{+2}$, $Mg^{+2}$, $Ni^{+2}$, $Zn^{+2}$.

I.a) 15min shoot (upregulated genes)

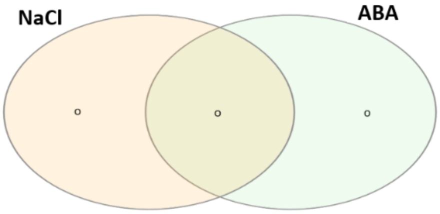


| Treatment | Number of genes | Name of genes |
|---|---|---|
| NaCl | 0 | - |
| ABA | 0 | - |
| NaCl+ABA | 0 | - |

I.b) 3h shoot (upregulated genes)

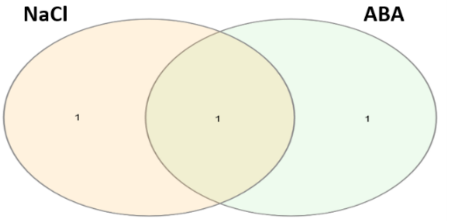


| Treatment | Number of genes | Name of genes |
|---|---|---|
| NaCl | 1 | OsGRAS19 |
| ABA | 1 | OsGRAS25 |
| NaCl+ABA | 1 | ΨOsGRAS5 |

I.c) 12h shoot (upregulated genes)

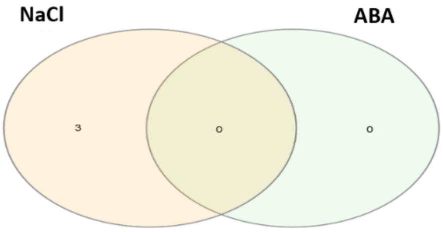


| Treatment | Number of genes | Name of genes |
|---|---|---|
| NaCl | 3 | OsGRAS12,ΨOsGRAS5, OsGRAS25 |
| ABA | 0 | - |
| NaCl+ABA | 0 | - |

I.d) 24h shoot (upregulated genes)

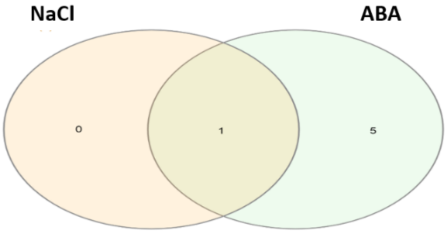


| Treatment | Number of genes | Name of genes |
|---|---|---|
| NaCl | 0 | |
| ABA | 5 | ΨOsGRAS2,OsGRAS12 OsGRAS24,OsGRAS25 OsSCR1 |
| NaCl+ABA | 1 | ΨOsGRAS5 |

I.e) 60h shoot (upregulated genes)

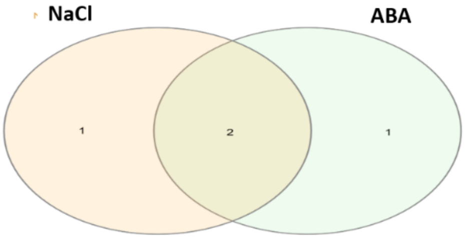


| Treatment | Number of genes | Name of genes |
|---|---|---|
| NaCl | 1 | OsGRAS12 |
| ABA | 1 | OsGRAS25 |
| NaCl+ABA | 2 | ΨOsGRAS5 OsGRAS24 |

II.a) 15min root (upregulated genes)

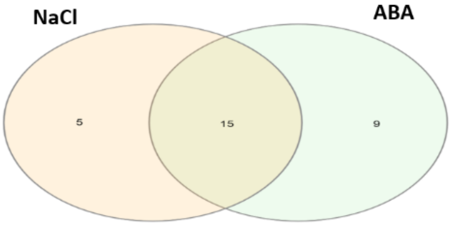


| Treatment | Number of genes | Name of genes |
|---|---|---|
| NaCl | 5 | OsGRAS8,OsGRAS11,OsGRAS19,OsGRAS22,OsGRAS26 |
| ABA | 9 | OsGRAS2,ΨOsGRAS2,ΨOsGRAS3,OsGRAS18,OsGRAS20,OsGRAS25,OsGRAS32,OsGRAS35,OsSCR1 |
| NaCl+ABA | 15 | OsGRAS5,OsGRAS10,OsGRAS12,OsSHR2,ΨOsGRAS4,OsGRAS15,OsSLR1,ΨOsGRAS5,OsGRAS24,OsCIGR1,OsSHR1,OsGRAS39,OsGRAS41,OsGRAS43,OsGRAS44 |

II.b) 3h root (upregulated genes)

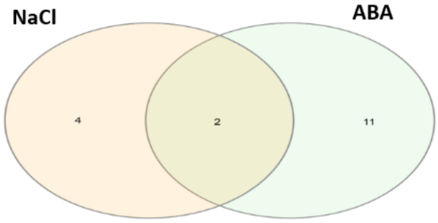


| Treatment | Number of genes | Name of genes |
|---|---|---|
| NaCl | 4 | OsGRAS10,ΨOsGRAS3,ΨOsGRAS4,OsSHR1 |
| ABA | 11 | OsGRAS5,ΨOsGRAS2,OsGRAS8,OsGRAS12,OsGRAS15,OsGRAS18,OsGRAS25,OsGRAS35,OsSCR1,OsGRAS41,OsGRAS44 |
| NaCl+ABA | 2 | OsGRAS24,OsGRAS39 |

II.c) 12h root (upregulated genes)

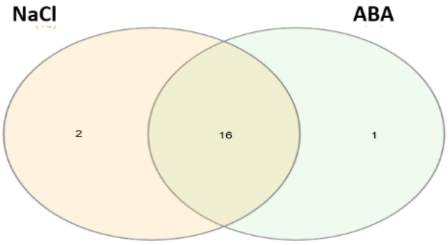


| Treatment | Number of genes | Name of genes |
|---|---|---|
| NaCl | 2 | OsSHR2,OsGRAS19 |
| ABA | 1 | OsGRAS41 |
| NaCl+ABA | 16 | OsGRAS2,OsGRAS5,ΨOsGRAS2,OsGRAS8,OsGRAS10,OsGRAS12,OsGRAS15,OsSLR1,OsGRAS22,OsGRAS24,OsGRAS25,OsSHR1,OsGRAS35,OsSCR1,OsGRAS39,OsGRAS44 |

II.d) 24h root (upregulated genes)

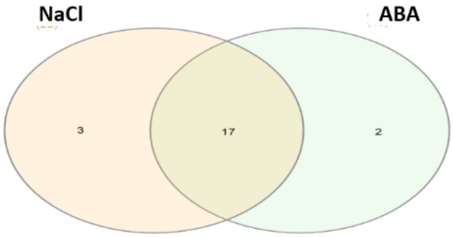


| Treatment | Number of genes | Name of genes |
|---|---|---|
| NaCl | 3 | OsGRAS11,OsSHR2,ΨOsGRAS9 |
| ABA | 2 | OsGRAS15,OsSHR1 |
| NaCl+ABA | 17 | OsGRAS2,OsGRAS5,ΨOsGRAS2,OsGRAS8,OsGRAS10,OsGRAS12,OsSLR1,OsGRAS19,OsGRAS22,OsGRAS24,OsGRAS25,OsGRAS35,OsSCR1,OsGRAS39,OsGRAS41,OsGRAS43,OsGRAS44 |

II.e) 60h root (upregulated genes)

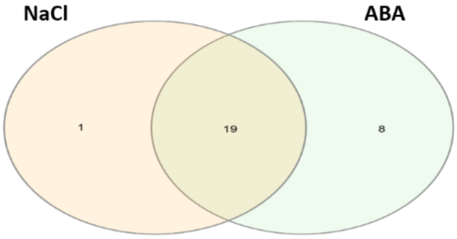


| Treatment | Number of genes | Name of genes |
|---|---|---|
| NaCl | 1 | ΨOsGRAS3 |
| ABA | 8 | OsGRAS2,OsGRAS3,OsGRAS18,OsGRAS20,OsGRAS22,OsGRAS41,ΨOsGRAS9,OsGRAS43 |
| NaCl+ABA | 19 | OsGRAS5,ΨOsGRAS2,OsGRAS8,OsGRAS10,OsGRAS11,OsGRAS12,OsSHR2,OsGRAS15,OsSLR1,OsGRAS19,ΨOsGRAS5,OsGRAS24,OsGRAS25,OsGRAS26,OsSHR1,OsGRAS35,OsSCR1,OsGRAS39,OsGRAS44 |

III.a) 15min shoot (downregulated genes)

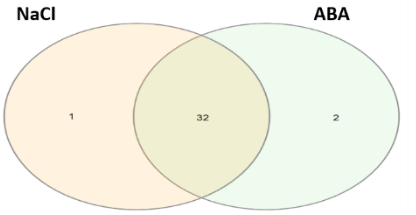


| Treatment | Number of genes | Name of genes |
| --- | --- | --- |
| NaCl | 1 | ΨOsGRAS2 |
| ABA | 2 | ΨOsGRAS5,OsCIGR1 |
| NaCl+ABA | 32 | OsGRAS8,OsGRAS10,OsGRAS11,OsGRAS12,OsSHR2, ΨOsGRAS3,ΨOsGRAS4,OsGRAS15,OsSLR1,OsGRAS18,OsGRAS19,OsGRAS20,OsGRAS22,OsGRAS23,OsGRAS24, OsGRAS25,OsGRAS26,OsGRAS28,OsGRAS32,OsCIGR2, OsSHR1,OsGRAS35,OsSCR1,ΨOsGRAS8,OsGRAS39, OsGRAS41,ΨOsGRAS9,OsGRAS43,OsGRAS44,OsGRAS47 ΨOsGRAS10,OsGRAS53 |

III.b) 3h shoot (downregulated genes)

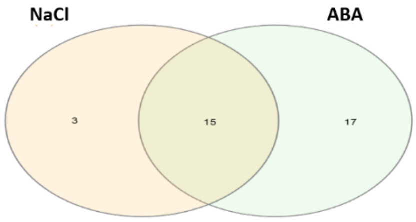


| Treatment | Number of genes | Name of genes |
| --- | --- | --- |
| NaCl | 3 | OsGRAS25,OsGRAS39,OsGRAS44 |
| ABA | 17 | OsGRAS2,OsGRAS3,OsGRAS5,OsGRAS7,OsGRAS8, OsGRAS10,OsGRAS11,OsSHR2,ΨOsGRAS3,ΨOsGRAS4, OsGRAS15,OsSLR1,OsGRAS18,OsGRAS20,OsGRAS22, OsGRAS24,OsCIGR1 |
| NaCl+ABA | 15 | OsGRAS23,OsGRAS26,OsGRAS28,OsGRAS32,OsCIGR2, OsSHR1,OsGRAS35,OsSCR1,ΨOsGRAS8,OsGRAS41, ΨOsGRAS9,OsGRAS43,OsGRAS47,ΨOsGRAS10,OsGRAS53 |

III.c) 12h shoot (downregulated genes)

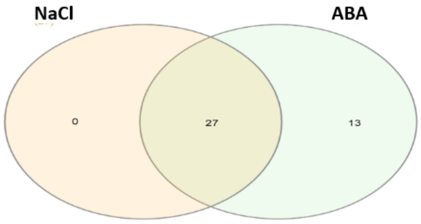


| Treatment | Number of genes | Name of genes |
| --- | --- | --- |
| NaCl | 0 | |
| ABA | 13 | OsGRAS1,OsGRAS2,OsGRAS3,OsGRAS5,OsGRAS12,OsGRAS19,ΨOsGRAS5,OsGRAS24,OsGRAS25, OsGRAS26,OsCIGR1,OsSCR1,ΨOsGRAS8 |
| NaCl+ABA | 27 | OsGRAS7,ΨOsGRAS2,OsGRAS8,OsGRAS10,OsGRAS11, OsSHR2,ΨOsGRAS3,ΨOsGRAS4,OsGRAS15,OsSLR1, OsGRAS18,OsGRAS20,OsGRAS22,OsGRAS23,OsGRAS28 ,OsGRAS32,OsCIGR2,OsSHR1,OsGRAS35,OsGRAS39,OsGRAS41,ΨOsGRAS9,OsGRAS43,OsGRAS44,OsGRAS47, ΨOsGRAS10,OsGRAS53 |

III.d) 24h shoot (downregulated genes)

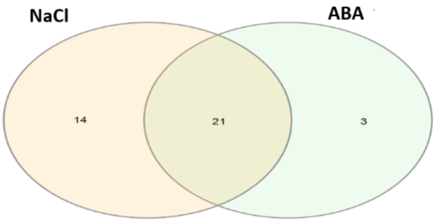


| Treatment | Number of genes | Name of genes |
| --- | --- | --- |
| NaCl | 14 | ΨOsGRAS2,OsGRAS8,OsGRAS10,OsGRAS11,OsGRAS12,OsSHR2,OsSLR1,OsGRAS19,OsGRAS24, OsGRAS25,OsGRAS26,OsSCR1,OsGRAS39,OsGRAS44 |
| ABA | 3 | OsGRAS2,OsGRAS3,OsGRAS5 |
| NaCl+ABA | 21 | OsGRAS7,ΨOsGRAS3,ΨOsGRAS4,OsGRAS15,OsGRAS18,OsGRAS20,OsGRAS22,OsGRAS23,OsGRAS28,OsCIGR1,OsGRAS32,OsCIGR2,OsSHR1, OsGRAS35,ΨOsGRAS8,OsGRAS41,ΨOsGRAS9, OsGRAS43,OsGRAS47,ΨOsGRAS10,OsGRAS53 |

III.e) 60h shoot (downregulated genes)

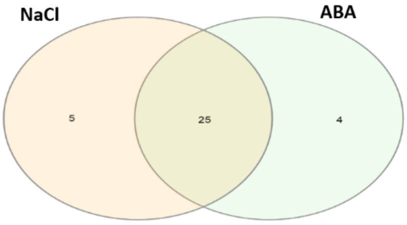


| Treatment | Number of genes | Name of genes |
| --- | --- | --- |
| NaCl | 5 | ΨOsGRAS2,ΨOsGRAS3,OsGRAS25,ΨOsGRAS8,OsGRAS44 |
| ABA | 4 | OsGRAS2,OsGRAS3,OsGRAS10,OsSLR1 |
| NaCl+ABA | 25 | OsGRAS7,OsGRAS8,OsGRAS11,OsSHR2,ΨOsGRAS4,OsGRAS15, OsGRAS18,OsGRAS19,OsGRAS20,OsGRAS22,OsGRAS23, OsGRAS26,OsGRAS28,OsCIGR1,OsGRAS32,OsCIGR2,OsSHR1, OsGRAS35,OsGRAS39,OsGRAS41,ΨOsGRAS9,OsGRAS43, OsGRAS47,ΨOsGRAS10,OsGRAS53 |

IV.a) 15min root (downregulated genes)

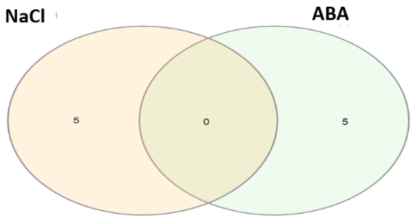


| Treatment | Number of genes | Name of genes |
|---|---|---|
| NaCl | 5 | ΨOsGRAS3,OsGRAS25,OsGRAS28 OsSCR1,ΨOsGRAS8 |
| ABA | 5 | OsGRAS1,OsGRAS3,OsGRAS7 OsGRAS8,OsGRAS19 |
| NaCl+ABA | 0 | |

IV.b) 3h root (downregulated genes)

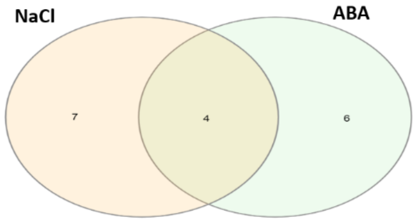


| Treatment | Number of genes | Name of genes |
|---|---|---|
| NaCl | 7 | OsGRAS1,OsGRAS3,OsGRAS11,ΨOsGRAS5, OsGRAS28,OsCIGR1,OsGRAS32 |
| ABA | 6 | OsGRAS7,OsSHR2,OsCIGR2,ΨOsGRAS8, OsGRAS43,ΨOsGRAS10 |
| NaCl+ABA | 4 | OsGRAS2,OsGRAS20,OsGRAS23,OsGRAS26 |

IV.c) 12h root (downregulated genes)

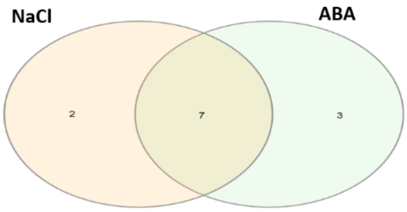


| Treatment | Number of genes | Name of genes |
|---|---|---|
| NaCl | 2 | OsCIGR1,ΨOsGRAS9 |
| ABA | 3 | ΨOsGRAS3,ΨOsGRAS4,OsGRAS28 |
| NaCl+ABA | 7 | OsGRAS7,OsGRAS23,OsGRAS32,OsCIGR2,Ψ OsGRAS8,ΨOsGRAS10,OsGRAS53 |

IV.d) 24h root (downregulated genes)

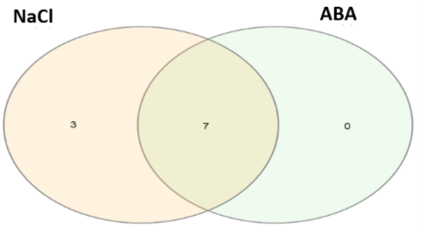


| Treatment | Number of genes | Name of genes |
|---|---|---|
| NaCl | 3 | OsGRAS7,ΨOsGRAS3,ΨOsGRAS4 |
| ABA | 0 | -- |
| NaCl+ABA | 7 | OsGRAS23,OsGRAS28,OsGRAS32,OsCIGR2, ΨOsGRAS8,ΨOsGRAS10,OsGRAS53 |

IV.d) 60h root (downregulated genes)

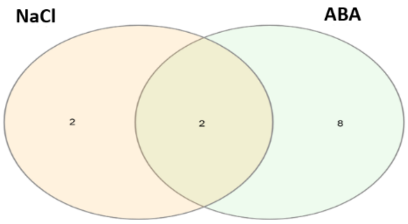


| Treatment | Number of genes | Name of genes |
|---|---|---|
| NaCl | 2 | OsCIGR1,ΨOsGRAS9 |
| ABA | 8 | ΨOsGRAS3,ΨOsGRAS4,OsGRAS23,OsGRAS28, OsGRAS32,OsCIGR2,ΨOsGRAS10,OsGRAS53 |
| NaCl+ABA | 2 | OsGRAS7,ΨOsGRAS8 |

Figure S4: Venn diagrams showing the number of genes upregulated in shoot (I.a-e) and root (II.a-e) or downregulated in shoot (III.a-e) and root (IV.a-e) during the course of treatment with NaCl and ABA. The corresponding number and list of genes under each treatment and in combination is mentioned in the figure.

Table S1: List of genes and their organization details

| Locus id | Gene name | Subfamily (according to Cenci and Rouard, 2017) | Chromosone number | Location | Gene size | Orientation | Splice forms | Introns | Exons |
|---|---|---|---|---|---|---|---|---|---|
| LOC_Os01g45860 | OsGRAS1 | OG-DELLA-2 | Chr1 | 26045843 - 26044166 | 1488 | 3'-5' | 1 | 0 | 1 |
| LOC_Os01g62460 | OsGRAS2 | OG-LISCL | Chr1 | 36158308 - 36161536 | 2463 | 5'-3' | 2 | 0 | 1 |
| LOC_Os01g65900 | OsGRAS3 | OG-PAT-3 | Chr1 | 38265509 - 38261836 | 1662 | 3'-5' | 1 | 2 | 3 |
| LOC_Os01g71970 | OsGRAS5 | OG-SCL3 | Chr1 | 41711881 - 41713811 | 1329 | 5'-3' | 1 | 0 | 1 |
| LOC_Os02g10360 | OsGRAS7 | OG-LS | Chr2 | 5453090 - 5451819 | 1272 | 3'-5' | 1 | 0 | 1 |
| LOC_Os02g21685 | ΨOsGRAS2 | OG-PAT-4 | Chr2 | 12888147 - 12887704 | 444 | 3'-5' | 1 | 0 | 1 |
| LOC_Os02g44360 | OsGRAS8 | OG-HAM-II | Chr2 | 26841585 - 26844331 | 2130 | 5'-3' | 1 | 0 | 1 |
| LOC_Os02g45760 | OsGRAS10 | OG-PAT-4 | Chr2 | 27856163 - 27853853 | 1857 | 3'-5' | 1 | 0 | 1 |
| LOC_Os03g09280 | OsGRAS11 | OG-PAT-1 | Chr3 | 4847121 - 4853832 | 1608 | 5'-3' | 3 | 0 | 1 |
| LOC_Os03g15680 | OsGRAS12 | OG-NSP2-1 | Chr3 | 8651497 - 8653224 | 1728 | 5'-3' | 1 | 0 | 1 |
| LOC_Os03g31880 | OsSHR2 | OG-SHR-1 | Chr3 | 18240730 - 18238256 | 1812 | 3'-5' | 1 | 0 | 1 |
| LOC_Os03g37900 | ΨOsGRAS3 | OG-DLT | Chr3 | 21050297 - 21050710 | 414 | 5'-3' | 1 | 0 | 1 |
| LOC_Os03g40080 | ΨOsGRAS4 | close to LISCL (unclassified) | Chr3 | 22262511 - 22267163 | 2334 | 5'-3' | 1 | 7 | 8 |
| LOC_Os03g48450 | OsGRAS15 | OG-LISCL | Chr3 | 27592832 - 27588776 | 2196 | 3'-5' | 4 | 0 | 1 |
| LOC_Os03g49990 | OsSLR1 | OG-DELLA-1 | Chr3 | 28512625 - 28515179 | 1878 | 5'-3' | 1 | 0 | 1 |
| LOC_Os03g51330 | OsGRAS18 | OG-SCL4/7 | Chr3 | 29370719 - 29373265 | 1737 | 5'-3' | 1 | 0 | 1 |
| LOC_Os04g35250 | OsGRAS19 | OG-SCLA | Chr4 | 21425721 - 21424207 | 1515 | 3'-5' | 1 | 0 | 1 |
| LOC_Os04g37440 | ΨOsGRAS5 | OG-DLT | Chr4 | 22312712 - 22312254 | 459 | 3'-5' | 1 | 0 | 1 |
| LOC_Os04g46860 | OsGRAS20 | OG-HAM-II | Chr4 | 27764666 - 27767328 | 2136 | 5'-3' | 1 | 0 | 1 |
| LOC_Os04g50060 | OsGRAS22 | OG-LISCL | Chr4 | 29857542 - 29860559 | 1911 | 5'-3' | 2 | 0 | 1 |
| LOC_Os05g31380 | OsGRAS23 | OG-SCL3 | Chr5 | 18234846 - 18236501 | 1656 | 5'-3' | 1 | 0 | 1 |
| LOC_Os05g31420 | OsGRAS24 | OG-SCL3 | Chr5 | 18275763 - 18277445 | 1683 | 5'-3' | 1 | 0 | 1 |
| LOC_Os05g40710 | OsGRAS25 | OG-SCR-3 | Chr5 | 23871516 - 23872997 | 1482 | 5'-3' | 1 | 0 | 1 |
| LOC_Os05g42130 | OsGRAS26 | OG-SCL32-2 | Chr5 | 24631172 - 24632715 | 1278 | 5'-3' | 1 | 0 | 1 |
| LOC_Os06g01620 | OsGRAS28 | OG-HAM-II | Chr6 | 365301 - 367066 | 1443 | 5'-3' | 1 | 0 | 1 |
| LOC_Os07g36170 | OsCIGR1 | OG-PAT-2 | Chr7 | 21616338 - 21620316 | 1716 | 5'-3' | 2 | 0 | 1 |
| LOC_Os07g38030 | OsGRAS32 | OG-SCR-2 | Chr7 | 22807706 - 22809384 | 1374 | 5'-3' | 1 | 0 | 1 |
| LOC_Os07g39470 | OsCIGR2 | OG-PAT-1 | Chr7 | 23650525 - 23654073 | 1635 | 5'-3' | 1 | 0 | 1 |
| LOC_Os07g39820 | OsSHR1 | OG-SHR-1 | Chr7 | 23868417 - 23871145 | 1809 | 5'-3' | 1 | 0 | 1 |
| LOC_Os07g40020 | OsGRAS35 | OG-SCL32-1 | Chr7 | 24014021 - 24018422 | 1422 | 5'-3' | 1 | 0 | 1 |
| LOC_Os11g03110 | OsSCR1 | OG-SCR-1 | Chr11 | 1119742 - 1123350 | 1956 | 5'-3' | 1 | 1 | 2 |
| LOC_Os11g04400 | ΨOsGRAS8 | OG-LISCL | Chr11 | 1829394 - 1832069 | 1650 | 5'-3' | 1 | 2 | 3 |
| LOC_Os11g04570 | OsGRAS39 | OG-SCL3 | Chr11 | 1939818 - 1933914 | 2565 | 5'-3' | 1 | 3 | 4 |
| LOC_Os11g06180 | OsGRAS41 | OG-NSP2-2 | Chr11 | 2950162 - 2952223 | 1419 | 5'-3' | 1 | 2 | 3 |
| LOC_Os11g11600 | ΨOsGRAS9 | close to LISCL (unclassified) | Chr11 | 6452428 - 6453736 | 1038 | 5'-3' | 1 | 4 | 5 |
| LOC_Os11g31100 | OsGRAS43 | OG-RAM1 | Chr11 | 18102879 - 18096831 | 2319 | 3'-5' | 1 | 2 | 3 |
| LOC_Os11g47870 | OsGRAS44 | OG-LISCL | Chr11 | 28870096 - 28872571 | 2079 | 5'-3' | 1 | 0 | 1 |
| LOC_Os11g47910 | OsGRAS47 | OG-LISCL | Chr11 | 28895187 - 28896974 | 1788 | 5'-3' | 1 | 0 | 1 |
| LOC_Os12g06540 | ΨOsGRAS10 | OG-NSP2-2 | Chr12 | 3170547 - 3172219 | 1389 | 5'-3' | 1 | 1 | 2 |
| LOC_Os12g38490 | OsGRAS53 | OG-LISCL | Chr12 | 23634672 - 23637678 | 2217 | 5'-3' | 1 | 0 | 1 |

Table S2: Chart depicting the protein properties of 40 GRAS genes under study. This includes their length, molecular weight, isoelectric points (*p*I), GRAVY indices, chelating ligands, low complexity region (LCR), localisation and secondary structure details

| NAME | LENGTH (aa) | MW (KDa) | *p*I | GRAVY index | Ligand residues | Metallic & non-metallic ligands | LCR | TargetP-2.0 prediction | Disordered % | α- helix % | β-sheet % |
|---|---|---|---|---|---|---|---|---|---|---|---|
| OsGRAS1 | 495 | 52067.35 | 5.06 | -0.017 | LEU168 HIS172 ASP193 PHE194 SER195 LEU196 MET197 GLN198 GLN201 ILE224 GLY225 PRO226 PRO229 SER262 LEU263 ASP264 VAL266 VAL284 GLN286 ARG289 LEU290 | Mg+2, SAM, Ca+2, SAH | Three (26-38,41-82,452-486) | - | 30 | 40 | 11 |
| OsGRAS2 | 820 | 90671.06 | 5.7 | -0.415 | LYS530 HIS534 ASP555 TYR556 GLY557 ILE558 TYR559 TYR560 GLN563 ILE586 ASP587 THR588 PRO589 GLN590 GLY592 SER624 ARG625 PHE626 GLU627 VAL629 MET647 LYS649 ASN652 | Mg+2, SAM, Ca+2, SAH | Four (85-97, 165-183,246-262,415-434) | - | 45 | 39 | 7 |
| OsGRAS3 | 553 | 61798.49 | 4.8 | -0.358 | MET260 PHE270 PHE274 ASP295 PHE296 ASP297 ILE298 ASN299 GLN300 GLN303 VAL326 ASP327 ASP328 ALA364 ASN365 ILE366 GLY367 VAL369 ALA387 GLN389 HIS392 | $Mg^{+2}$, SAM, NAP, SAH | Two (91-107,183-553) | - | 35 | 39 | 10 |
| OsGRAS5 | 442 | 48023.77 | 6.15 | -0.079 | LEU139 ASP164 LEU165 GLY166 GLY167 ALA168 ASP169 GLN172 VAL195 HIS196 GLU197 ARG228 LEU229 ASP230 SER250 GLN252 | $Mg^{+2}$, SAM, $Ca^{+2}$, SAH | Four (46-73,89-105,260-270,367-383) | - | 24 | 45 | 12 |
| OsGRAS7 | 423 | 44138.74 | 5.56 | -0.113 | LEU122 HIS126 ASP153 LEU154 ASP155 ALA156 ALA157 HIS158 GLN161 ALA188 GLY189 THR190 LEU226 ALA227 VAL247 PHE249 LYS252 | Mg+2, SAM, Ca+2, SAH | Two (21-29,39-420) | - | 22 | 44 | 12 |
| ΨOsGRAS2 | 147 | 15305.11 | 9.18 | -0.128 | | | Three (38-62,81-91,131-144) | - | 48 | 58 | 0 |
| OsGRAS8 | 709 | 74248.23 | 5.65 | -0.02 | LEU434 ASP461 PHE462 ASP463 GLY465 VAL466 GLN469 PHE496 MET497 LEU532 ASP533 ALA534 PHE535 PRO554 | Mg+2, SAM, Ca+2, SAH | Five (47-85,88-106,153-165,195-217,258-273) | - | 51 | 31 | 8 |
| OsGRAS10 | 618 | 64200.52 | 9.27 | -0.115 | PHE335 ASP360 PHE361 ASP362 VAL363 SER364 GLN367 VAL390 ALA391 ASP392 CYS430 ARG431 ALA432 PRO433 ILE435 ALA453 THR455 ARG458 | SAM, Mg+2, Ca+2,SAH | Six (23-35,40-50,62-72,98-116,143-160,162-179) | - | 38 | 45 | 9 |
| OsGRAS11 | 535 | 59647.91 | 5.86 | -0.375 | MET242 PHE252 TYR256 ASP277 PHE278 GLN279 ALA281 GLN282 ILE308 ASP309 ASP310 HIS315 ALA346 ALA347 SER348 HIS349 VAL351 ALA369 TYR370 GLN371 HIS374 | $Mg^{+2}$, SAM, $Ca^{+2}$, SAH | One (80-93) | - | 35 | 40 | 10 |
| OsGRAS12 | 575 | 60848.2 | 5.04 | -0.106 | MET231 HIS235 ASP256 TYR257 ASP258 ILE259 ALA260 GLU261 GLN264 VAL289 SER290 ARG291 GLY294 GLY295 LEU325 ASP328 VAL349 LEU350 HIS351 | Mg+2, SAM, Ca+2, SAH | Three (15-34,55-65,100-116) | - | 35 | 43 | 9 |
| OsSHR2 | 603 | 64247.77 | 5.93 | -0.354 | THR278 HIS282 LEU323 SER324 ASN325 THR326 PHE327 THR329 VAL354 VAL355 PRO356 THR357 HIS393 GLY395 ASP396 LEU397 VAL421 ASN422 | SAM, SAH, NAD, ATP | Four (11-18,31-94,118-141,162-185) | - | 41 | 36 | 9 |
| ΨOsGRAS3 | 137 | 15192.7 | 7.87 | -0.07 | | | -- | - | 31 | 71 | 0 |
| ΨOsGRAS4 | 777 | 88331.49 | 5.53 | -0.486 | | | -- | - | 40 | 34 | 9 |
| OsGRAS15 | 731 | 82218.82 | 6.44 | -0.483 | LYS443 HIS447 ASP468 PHE469 GLY470 ILE471 TYR472 PHE473 ILE499 ASP500 VAL501 PRO502 GLN503 GLY505 PRO508 LYS538 TRP539 GLU540 ILE542 LEU560 ARG562 ASN565 | Mg+2, SAM, Ca+2, SAH | -- | - | 40 | 39 | 8 |
| OsSLR1 | 625 | 65406.24 | 5.14 | -0.111 | LEU323 HIS327 ASP348 PHE349 GLY350 ILE351 LYS352 GLN353 GLN356 VAL379 GLY380 PRO381 PRO382 GLN383 PRO384 ASP385 THR417 LEU418 ALA419 VAL446 PHE447 GLU448 ARG451 | SAM, SAH | Four (9-17,128-140,185-207,209-232) | - | 40 | 44 | 9 |

| | | | | | | | | | | | |
|---|---|---|---|---|---|---|---|---|---|---|---|
| OsGRAS18 | 578 | 62477.53 | 5.63 | -0.209 | TYR283 HIS297 ASP318 PHE319 GLY320 ILE321 VAL322 VAL350 PRO351 SER352 PRO353 LEU354 LEU355 VAL390 MET411 LEU412 GLN413 TYR415 HIS416 | SAM, Ca+2,SAH | One (144-206) | - | 36 | 44 | 10 |
| OsGRAS19 | 504 | 52579.87 | 8.67 | -0.224 | TYR169 TYR175 HIS179 ASP202 PHE203 ASP204 VAL205 SER206 TYR207 GLN210 PHE252 GLY253 ALA254 ASN285 ASN286 GLY287 SER288 THR290 VAL308 | Mg+2, SAM, SAH | One (10-42) | - | 31 | 45 | 11 |
| ΨOsGRAS5 | 152 | 16672.18 | 9.43 | -0.472 | | | Two (104-124,130-146) | - | 38 | 38 | 14 |
| OsGRAS20 | 711 | 74007.88 | 5.57 | 0.004 | LEU437 ASP464 PHE465 ASP466 LEU467 GLY468 VAL469 GLN472 PHE499 VAL500 SER501 LEU535 ASP536 ALA537 PRO557 VAL558 | SAM, NAP, Ca+2, SAH | Eight (52-70,79-92,98-114,185-215,221-234,243-255,277-286,302-312) | - | 52 | 32 | 8 |
| OsGRAS22 | 636 | 71655.98 | 5.51 | -0.424 | PHE338 LYS348 ASP373 TYR374 GLY375 ILE376 GLN377 TYR378 VAL404 ASP405 LEU406 ARG412 ALA442 LYS443 TRP444 ASP445 ILE447 LEU467 ARG469 ASN472 MET473 | Mg+2, SAM, SAH | Three (2-15,56-76,237-251) | - | 37 | 38 | 8 |
| OsGRAS23 | 551 | 56280.97 | 5.41 | 0.197 | VAL182 ASP208 LEU209 GLY210 GLY211 GLY212 VAL213 ASP214 GLN217 VAL241 ASN242 GLU243 SER274 ILE275 GLU276 ASN303 GLN305 ARG308 | Mg+2, SAM, Ca+2, SAH | Three (6-27,51-59,513-541) | - | 36 | 41 | 10 |
| OsGRAS24 | 560 | 58183.56 | 7.85 | 0.182 | LEU248 ASP273 LEU274 GLY275 GLY276 ILE277 HIS278 HIS281 VAL305 HIS306 GLU307 SER338 VAL339 GLU340 THR360 GLN362 ARG365 | Mg+2,SAM, SAH | Four (5-20,32-52,55-83,89-141) | - | 42 | 36 | 9 |
| OsGRAS25 | 493 | 51644.41 | 5.75 | 0.008 | TYR207 ASP228 LEU229 ASP230 VAL231 VAL232 PRO233 GLY234 GLN238 PHE261 GLY262 MET263 ARG294 PRO295 GLY296 ALA298 TRP317 LEU318 ARG319 HIS320 | SAM, SAH | Two (13-36,98-111) | - | 30 | 45 | 10 |
| OsGRAS26 | 425 | 45430.82 | 5.81 | 0.017 | HIS131 PHE135 ASP156 LEU157 SER158 VAL159 THR160 HIS161 GLN164 PRO187 SER188 VAL189 ARG190 PRO191 ALA192 SER226 ALA227 THR228 THR229 GLN266 SER267 TRP268 | SAM, Mg+2,SAH | One (9-20) | - | 21 | 50 | 12 |
| OsGRAS28 | 480 | 51017.01 | 5.46 | -0.058 | VAL213 ASP238 PHE239 ASP240 VAL241 GLY242 PHE243 LEU273 VAL274 SER275 PRO276 GLY277 SER278 SER279 PHE309 THR334 | Mg+2, SAM, Ca+2, SAH | One (35-66) | - | 36 | 45 | 11 |
| OsCIGR1 | 571 | 64602.19 | 5.87 | -0.478 | MET278 PHE288 TYR292 ASP313 PHE314 GLN315 ILE316 ALA317 GLN318 GLN321 ILE344 ASP345 ASP346 VAL382 TYR383 ALA384 THR385 VAL387 THR405 GLN407 | Mg+2, SAM, Ca+2, SAH | -- | - | 40 | 36 | 9 |
| OsGRAS32 | 457 | 49096.59 | 5.51 | -0.127 | PHE148 ASN152 VAL158 HIS162 ASP183 LEU184 ASP185 ILE186 MET187 GLN188 GLN191 LEU214 GLY215 ALA216 LYS247 ILE248 GLY249 VAL251 VAL269 TRP271 HIS274 | Mg+2, SAM, Ca+2, SAH | Three (8-45,55-62,437-449) | - | 27 | 47 | 14 |
| OsCIGR2 | 544 | 60108.16 | 6 | -0.324 | MET251 PHE261 TYR265 ASP286 PHE287 HIS288 ILE289 SER290 GLN291 GLN294 ILE317 ASP318 ASP319 ILE355 SER356 GLY357 SER358 VAL360 THR378 GLU380 HIS383 ILE384 | SAM, Mg+2, SAH | -- | - | 35 | 40 | 10 |
| OsSHR1 | 602 | 64709.33 | 5.61 | -0.4 | HIS286 LEU316 SER317 ASN318 THR319 PHE320 THR322 VAL347 VAL348 SER349 ALA350 HIS388 GLY390 ASP391 LEU392 VAL416 ASN417 MET527 | SAM, Ca+2,SAH, Zn+2 | Three (11-40,46-80,122-147) | - | 39 | 40 | 7 |
| OsGRAS35 | 473 | 50596.45 | 5.31 | 0.022 | TYR174 LEU196 SER197 THR198 THR199 HIS200 ALA226 ASP227 VAL228 ALA229 TYR241 THR266 SER274 LEU275 VAL276 HIS297 MET298 LEU299 THR302 | Mg+2, SAM, SAH | Two (9-25,34-54) | - | 27 | 43 | 11 |

| | | | | | | | | | | | |
|---|---|---|---|---|---|---|---|---|---|---|---|
| OsSCR1 | 651 | 69918.23 | 5.91 | -0.221 | PHE364 HIS378 ASP399 LEU400 ASP401 ILE402 MET403 GLN407 LEU430 GLY431 ALA432 ASP462 LYS463 ALA464 LEU485 HIS487 LEU489 TYR490 ASP491 VAL492 THR493 | $Ni^{+2,}$ $Mg^{+2}$, SAM, NAP, SAH | Six (3-55,86-101,115-134,148-157,188-229,235-279) | - | 48 | 37 | 9 |
| ΨOsGRAS8 | 549 | 59950.36 | 4.98 | -0.299 | | | Four (29-40,108-122,140-147,177-193) | - | 34 | 48 | 8 |
| OsGRAS39 | 854 | 94267.73 | 7.45 | -0.09 | LEU495 TYR499 ASP520 PHE521 SER522 GLY523 PRO524 ALA525 ALA526 ASN527 GLN530 VAL553 HIS554 ASP555 ALA585 LYS586 LEU587 ASP588 VAL612 GLN614 ARG617 | SAM, SAH, $Ca^{+2}$ | One (370-391) | Chloroplast | 23 | 45 | 12 |
| OsGRAS41 | 472 | 52312.55 | 4.78 | 0.111 | VAL135 HIS139 ASP160 LEU161 ASN162 ILE163 GLY164 GLU165 GLN168 ILE189 THR190 THR191 VAL225 HIS226 ASN227 GLU228 GLU230 THR248 THR249 SER250 | Mg+2, SAM, Ca+2, SAH | -- | - | 23 | 53 | 7 |
| ΨOsGRAS9 | 345 | 37286.47 | 10.13 | -0.552 | | | One (216-254) | - | 35 | 49 | 8 |
| OsGRAS43 | 772 | 81094.36 | 6.18 | -0.156 | | | Eight (2-13,165-177,183-201,223-230,280-288,293-306,326-341,355-364) | - | 39 | 36 | 11 |
| OsGRAS44 | 692 | 77442.34 | 5.01 | -0.394 | | | Four (14-27,82-111,209-218,279-299) | - | 39 | 43 | 8 |
| OsGRAS47 | 595 | 66666.1 | 5.99 | -0.372 | | | One (201-211) | - | 33 | 48 | 10 |
| ΨOsGRAS10 | 462 | 50587.65 | 4.53 | -0.137 | | | Two (22-30,47-60) | - | 44 | 44 | 7 |
| OsGRAS53 | 738 | 81365.04 | 5.17 | -0.305 | | | Nine (53-65,116-135,150-162,241-260,337-356,796-808,859-878,893-905,984-1003) | - | 39 | 40 | 7 |

Table S3: List of domains observed in GRAS genes. The number, position and sequences of the observed domains and LCR are provided in the chart.

| Name | Domain | Position | Sequence | LCR | Position | Sequence |
|---|---|---|---|---|---|---|
| OsGRAS1 | GRAS | 86-449 | LVHLLMSCAGAIEAGDHALASAQLADSHAALAAVSAASGIGRVAVHFTTALSRRLFPSPVAPPTTDAEHAFLYHHFYEACPYLKFAHFTANQAILEAFHGCDHVHVIDFSLMQGLQWPALIQALALRPGGPPFLRITGIGPPSPTGRDELRDVGLRLADLARSVRVRFSFRGVAANSLDEVRPWMLQIAPGEAVAFNSVLQLHRLLGDPADQAPIDAVLDCVASVRPKIFTVIEQEADHNKTGFLDRFTEALFYYSAVFDSLDAASASGGAGNAMAEAYLQREICDIVCGEGAARRERHEPLSRWRDRLTRAGLSAVPLGSNALRQARMLVGLFSGEGHSVEEADGCLTLGWHGRPLFSASAWE | LCR1 | 26-38 | PPPAAVAPDDGVG |
| | | | | LCR2 | 41-82 | DPPAGADVDAAALPEFAAAFPPCAPDAAAAVLAMRREEEEVA |
| | | | | LCR3 | 452-486 | GDGGGDNNNNSNSNVSGSSGSDSNNSGSSNGKSSG |
| OsGRAS2 | GRAS | 443-813 | LETLLIHCAQSVATDDRRSATELLKQIRQHAHANGDGDQRLAHCFANGLEARLAGTGSQIYKNYTITRLPCTDVLKAYQLYLAACPFKKISHYFANQTILNAVEKAKKVHIVDYGIYYGFQWPCLIQRLSNRPGGPPKLRITGIDTPQPGFRPAERTEETGRYLSDYAQTFNVPFEFQAIASRFEAVRMEDLHIEEDEVLIVNCMFKFKNLMDESVVAESPRNMALKTIRKMNPHVFIHGVVNGSYNAPFFVTRFREALFHYSAIFDMLETNIPKDNEQRLLIESALFSREAINVISCEGLERMERPETYKQWQVRNQRVGFKQLPLNQDMMKRAREKVRCYHKDFIIDEDNRWLLQGWKGRILFALSTWK | LCR1 | 85-97 | VSVSASAASSAAA |
| | | | | LCR2 | 165-183 | SSAANSCNSLSPCNCSSSS |
| | | | | LCR3 | 246-262 | SQSSSFASSNGSSVTFS |
| | | | | LCR4 | 415-434 | KHSGGGHGKGSSHGKGRGKK |
| OsGRAS3 | GRAS | 183-553 | PKQLLFDCAMALSDYNVDEAQAIITDLRQMVSIQGDPSQRIAAYLVEGLAARIVASGKGIYKALSCKEPPTLYQLSAMQILFEICPCFRFGFMAANYAILEACKGEDRVHIIDFDINQGSQYITLIQFLKNNANKPRHLRITGVDDPETVQRTVGGLKVIGQRLEKLAEDCGISFEFRAVGANIGDVTPAMLDCCPGEALVVNFAFQLHHLPDESVSIMNERDQLLRMVKGLQPKLVTLVEQDANTNTAPFQTRFREVYDYYAALFDSLDATLPRESPDRMNVERQCLAREIVNILACEGPDRVERYEVAGKWRARMTMAGFTPCPFSSNVISGIRSLLKSYCDRYKFEEDHGGLHFGWGEKTLIVSSAWQ | LCR | 91-107 | SNVSQQNSQSISDNQSS |
| OsGRAS5 | GRAS | 50-437 | LIHLLLNCAAAAAAGRLDAANAALEHIASLAAPDGDAMQRVAAAFAEALARRALRAWPGLCRALLLPRASPTPAEVAAARRHFLDLCPFLRLAGAAANQSILEAMESEKIVHVIDLGGADATQWLELLHLLAARPEGPPHLRLTSVHEHKELLTQTAMALTKEAERLDVPFQFNPVVSRLDALDVESLRVKTGEALAICSSLQLHCLLASDDDAAAVAGGDKERRSPESGLSPSTSRADAFLGALWGLSPKVMVVAEQEASHNAAGLTERFVEALNYYAALFDCLEVGAARGSVERARVERWLLGEEIKNIVACDGGERRERHERLERWARRLEGAGFGRVPLSYYALLQARRVAQGLGCDGFKVREEKGNFFLCWQDRALFSVSAWR | | | |
| OsGRAS7 | GRAS | 39-420 | ARGLVLACADLVHRGDLDGARRVAEAVLAAADPRGEAGDRLAHHFARALLALRGGGKGGHGGGGGGVVPSSAAYLAYIKIAPFLRFAHLTANQAILEAAAADAGGAHRRVLHIVDLDAAHGVQWPPLLQAIADRADPAVGPPPEVRLTGAGTDRDVLLRTGDRLRAFSSSLNLPFRFHPLILPCTAELAADPTAALELHPDETLAVNCVLFLHKLGGDGELAAFLRWVKSMNPAVVTIAEREGVLGGDVDDDNVPDELPRRVAAAMDYYSSVFDALEATVPPASADRLAVEQEILSREIDAAVAAPGAGGGGRARDFDAWASAARAAGLAPRPLSAFAASQARLLLRLHYPSEGYKADDDGGRGACFLRWQQRPLMSVSSWQ | LCR | 21-29 | QPQPQPQPP |
| ΨOsGRAS2 | GRAS | 62-138 | SRQLLSEAAAAIANGNHIVAASLLSALKLSVNPQGDAEQRLVAMMVAALSSCVGTSPSQHLADLYIGVGRRRWSEDR | | | |
| OsGRAS8 | GRAS | 349-708 | LLDELAAAAKATEAGNSVGAREILARLNQQLPQLGKPFLRSASYLKEALLLALADSHHGSSGVTSPLDVALKLAAYKSFSDLSPVLQFTNFTATQALLDEIGGMATSCIHVIDFDLGVGGQWASFLQELAHRRGAGGMALPLLKLTAFMSTASHHPLELHLTQDNLSQFAAELRIPFEFNAVSLDAFNPAELISSSGDEVVAVSLPVGCSARAPPLPAILRLVKQLCPKVVVAIDHGGDRADLPFSQHFLNCFQSCVFLLDSLDAAGIDADSACKIERFLIQPRVEDAVIGRHKAQKAIAWRSVFAATGFKPVQLSNLAEAQADCLLKRVQVRGFHVEKRGAALTLYWQRGELVSISSWR | LCR1 | 47-85 | SPSPPYSTSTLSSSLGGGSADSTGVAAVSESSTAAAGAT |
| | | | | LCR2 | 88-106 | GAPGEHGGGGKEEWGGGCE |
| | | | | LCR3 | 153-165 | QPGPPLPVLQQPL |
| | | | | LCR4 | 195-217 | SSSGAHTATGGGGKASLGFGLFS |
| | | | | LCR5 | 258-273 | PPNPAAALFMPLPPFP |
| OsGRAS10 | GRAS | 253-618 | SRQLLSEAAAAVADGNHTAAASLLSALKLSANPRGDAEQRLVAMMVAALSSRVGTGPSQHLADLYSGEHRAACQLLQDVSPCFGLALHGANLAILDAVAGHRAIHLVDFDVSAAQHVALIKALADRRVPATSLKVTVVADPTSPFTPAMTQSLAATCERLKKLAQQAGIDFRFRAVSCRAPEIEASKLGCEPGEALAVNLAFTLSRVPDESVSPANPRDELLRRVRALGPRVVTLVEQELNTNTAPMAARFSDASAHYGAVLESLDATLGRDSADRTRAEAALASKVANAVGREGPDRVERCEVFGKWRARFGMAGFRAVAIGEDIGGRVRARLGPALPAFDVKLDNGRLGVGWMGRVVTVASAWR | LCR1 | 23-35 | AALQAAARQQSQQ |
| | | | | LCR2 | 40-50 | GAGGAGVTGGV |
| | | | | LCR3 | 62-72 | QQQRQVAAQQA |
| | | | | LCR4 | 98-116 | GISGLSSGFGGISQQQPSS |
| | | | | LCR5 | 143-160 | TAQNQAVARAPAARPATA |
| | | | | LCR6 | 162-179 | ELVLLQELEKQLLGDDEE |
| OsGRAS11 | GRAS | 166-535 | LKQVIAACGKAVDENSWYRDLLISELRNMVSISGEPMQRLGAYMLEGLVARLSSTGHALYKSLKCKEPTSFELMSYMHLLYEICPFFKFGYMSANGAIAEAVKGENFVHIIDFQIAQGSQWATMIQALAARPGGPPYLRITGIDDSNSAHARGGGLDIVGRRLFNIAQSCGLPFEFNAVPAASHEVMLEHLDIRSGEVIVVNFAYQLHHTPDESVGIENHRDRILRMVKGLSPRVVTLVEQEANTNTAPFFNRYLETLDYYTAMFEAIDVACPRDDKKRISTEQHCVARDIVNLIACEGAERVERHEPFGKWRARLSMAGFRPYPLSALVNNTIKKLLDSYHSYYKLEERDGALYLGWKNRKLVVSSAWR | LCR | 80-93 | HSSTSSHISGSPIS |
| OsGRAS12 | GRAS | 132-526 | LLHLLMAAAEALSGPHKSRELARVILVRLKEMVSHTASANAAASNMERLAAHFTDALQGLLDGSHPVGGSGRQAAAAASHHHAGDVLTAFQMLQDMSPYMKFGHFTANQAILEAVSGDRRVHIVDYDIAEGIQWASLMQAMTSRADGVPAPHLRITAVSRSGGGGARAVQEAGRRLSAFAASIGQPFSFGQCRLDSDERFRPATVRMVKGEALVANCVLHQAAATTTIRRPTGSVASFLSGMAALGAKLVTVVEEEGEAEKDDDGDSAGDAAAGGFVRQFMEELHRYSAVWDSLEAGFPTQSRVRGLVERVILAPNIAGAVSRAYRGVDGEGRCGWGQWMRGSGFTAVPLSCFNHSQARLLLGLFNDGYTVEETGPNKIVLGWKARRLMSASVWA | LCR1 | 15-34 | SGCGSTTTTSSASSLDDGTG |
| | | | | LCR2 | 55-65 | DDDGGHDLHGL |
| | | | | LCR3 | 100-116 | GNGSNPSSTTTTNPGSP |
| OsSHR2 | GRAS | 188-502 | AAQLLMECARAVAGRDSQRVQQLMWMLNELASPYGDVDQKLASYFLQGLFARLTTSGPRTLRTLATASDRNASFDSTRRTALKFQELSPWTPFGHVAANGAILESFLEAAAAGAAASSSSSSSSSTPPTRLHILDLSNTFCTQWPTLLEALATRSSDDTPHLSITTVVPTAAPSAAAQRVMREIGQRLEKFARLMGVPFSFRAVHHSGDLADLDLAALDLREGGATAALAVNCVNALRGVARGRDAFVASLRRLEPRVVTVVEEEADLAAPEADASSEADTDAAFVKVFGEGLRFFSAYMDSLEESFPKTSNERLSLERAVGRAIVDLVSCPASQSAERRETAASWARRMRSAGFSPAAFSEDVADDVRSLLRRYKEGWSMRDAGGATDDAAGAAAAGAFLAWKEQPVVWASAWK | LCR1 | 11-18 | HHHHHHQH |
| | | | | LCR2 | 31-94 | SYPSSRGSTSSPSSHHTHNHTYYHHSHSHYNNNSNTNYYYQGGGGGGGGYYYAEEQQPAAYLEE |
| | | | | LCR3 | 118-141 | SGTGAPSSAPVPPPPSATTSSAGG |
| | | | | LCR4 | 162-185 | GGSPAVPSSSGAGAGAGAAPSSSG |
| ΨOsGRAS3 | GRAS | 38-137 | LVRMLTACADSVSAGNHEAAIYYLARLCEMASLAGPMPIHRVAAYFIEVLTLRVVRMWPHMFNISPPRELTNDAFSGDDDAMALRILNTITPILLLGKHS | | | |
| ΨOsGRAS4 | GRAS | 360-435 | LRTLLINCAQAVSVSNHSLASDILKIIRHHASPTGDDSQRLALCLAYCLDVRLTGTGSQIYHKFITKRRNVKDILK | | | |
| | GRAS | 428-672 | RNVKDILKVHIIDFGICFGFQWPSLFEELAKIEDGPPKLRITGIELPESGFRPYARSNNIGLRLADYAKTFNIPFEYQHISSNKWEALSPEDFNIEKDEVLIVNCIYRIKDLGDETISINSARSRVLNTIRMMKPKVFVQGVLNGSYGVPFFLTRFKEVMYHYNSLFDMLDKNIPRDNETRMIIERDIYQYIMLNVIACEGPERIERPESYKKWKVRNLKAGLVQLPLNPAIVRETQDMSSDKAS | | | |
| | RPT1 | 730-756 | VSDDFYHVSGDTREISCDTYQVLDDFY | | | |
| | RPT1 | 751-777 | VLDDFYHVSGDTYEISDDTYRISGDSY | | | |
| OsGRAS15 | GRAS | 356-727 | LRTLLIHCAQAVAADDRRTANELLKQIRQHAKPNGDGSQRLAYCFADGLEARLAGTGSQLYHKLVAKRTTASDMLKAYHLYLAACPFKRLSHFLSNQTILSLTKNASKVHIIDFGIYFGFQWPCLIRRLFKREGGPPKLRITGIDVPQPGFRPTERIEETGQRLAEYAEKIGVPFEYQGIASKWETICVEDLNIKKDEVVIVNCLYRFRNLIDETVAIDSPRNRVLNTIRQVNPAIFIHGIVNGSYSVPFFITRFREALFHFSALFDMLETTVPRDDAQRALIERDLFGREALNVIACEGSDRVERPETYKQWQVRNLRAGFVQSPLNQDIVLKAKDKVKDIYHKDFVIDEDSEWLLQGWKGRIIYAISTWK | LCR1 | 50-64 | SSAASSTASRAAVSS |
| | | | | LCR2 | 133-148 | PLDSPSESSTSSYPHS |
| OsSLR1 | DELLA | 39-120 | DELLAALGYKVRSSDMADVAQKLEQLEMAMGMGGVSAPGAADDGFVSHLATDTVHYNPSDLSSWVESMLSELNAPLPPIPPA | LCR1 | 9-17 | GGSSGGGSS |
| | GRAS | 241-621 | LVHALLACAEAVQQENFAAAEALVKQIPTLAASQGGAMRKVAAYFGEALARRVYRFRPADSTLLDAAFADLLHAHFYESCPYLKFAHFTANQAILEAFAGCHRVHVVDFGIKQGMQWPALLQALALRPGGPPSFRLTGVGPPQPDETDALQQVGWKLAQFAHTIRVDFQYRGLVAATLADLEPFMLQPEGEADANEEPEVIAVNSVFELHRLLAQPGALEKVLGTVHAVRPRIVTVVEQEANHNSGSFLDRFTESLHYYSTMFDSLEGGSSGQAELSPPAAGGGGGTDQVMSEVYLGRQICNVVACEGAERTERHETLGQWRNRLGRAGFEPVHLGSNAYKQASTLLALFAGGDGYRVEEKEGCLTLGWHTRPLIATSAWR | LCR2 | 128-140 | STSSTVTGGGGSG |
| | | | | LCR3 | 185-207 | GGGSTSSSSSSSSLGGGASRGS |
| | | | | LCR4 | 209-232 | VEAAPPATQGAAAANAPAVPVVVV |
| OsGRAS18 | SCOP | 15-95 | QQVIQQQQQQQQQQRHHHHHHLPPPPPPQSMAPHHHQQKHHHHHQQMPAMPQAPPSSHGQIPGQLAYGGGAAWPAGEHF | LCR1 | 144-206 | TTPPPPVPSPPPTHAAATATATAATAAPRPEAAPALLPQPAAATPVACSSPSPSSADASCSAP |
| | GRAS | 207-578 | ILQSLLSCSRAAATDPGLAAAELASVRAAATDAGDPSERLAFYFADALSRRLACGTGAPPSAEPDARFASDELTLCYKTLNDACPYSKFAHLTANQAILEATGAATKIHIVDFGIVQGIQWAALLQALATRPEGKPTRIRITGVPSPLLGPQPAASLAAT | | | |

| | | | | | | |
|---|---|---|---|---|---|---|
| | | | NTRLRDFAKLLGVDFEFVPLLRPVHELNKSDFLVEPDEAVAVNFMLQLYHLLGDSDELVRRVLRLAKSLSPAVVTLGEYEVSLNRAGFVDRFANALSYYRSLFESLDVAMTRDSPERVRVERWMFGERIQRAVGPEEGADRTERMAGSSEWQTLMEWCGFEPVPLSNYARSQADLLLWNYDSKYKYSLVELPPAFLSLAWEKRPLLTVSAWR | | | |
| OsGRAS19 | GRAS | 86-495 | LVRLLLSAVAAGEAGDARAAAAALREVDRRASCRGGGDPAQRVAACYAAALAPRLAAGLRPARSSPAAPAAARAEQFLAYTMFYQASPFYQFAHFTANQAIVEAFESGGRRRLHVVDFDVSYGFQWPSLIQSLSDAAAAATSSSSHDDDDNGGGCGDGPVSLRITGFGASADELRETEARLRRFAAGCPNLRFEFEGILNNGSNTRHDCTRIDDDATVVVNLVFPASSREACAATRMAYINSLNPSMVFLIEKHDGGGGLTGGDNTTTGRSASLLPRFAANLRYFAAVFDSLHECLPADSAERLAIERDHLGREIADAVASLDHQHRRRHGGGGGGGDHAAASWNWKAAMEGAGLDGVKLSSRTVSQAKLLLKMKSGCGGGGFRVVEGDGGMAMSLAWRDMALATATLWR | LCR | 10-42 | DGGGGGDAAAAVAKKSKVVGGGAVVVDGVGSSA |
| ΨOsGRAS5 | GRAS | 1-80 | MHYLRYYDAAFDAVDAAGLLETRPARAKVEEMFAREIRNAVAFEGAERFERHESFAGRRRRMEDGGGLQWGSKAEEKCLL | LCR1 | 104-124 | SLPPAVAAAPLVLPLPRASAA |
| | | | | LCR2 | 130-146 | APMPPTAAPLVLPPPLP |
| OsGRAS20 | GRAS | 352-710 | LLDELAAAAKATEVGNSIGAREILARLNQQLPPIGKPFLRSASYLKDALLLALADGHHAATRLTSPLDVALKLTAYKSFSDLSPVLQFANFTVTQALLDEIASTTASCIRVIDFDLGVGGQWASFLQELAHRCGSGGVSLPMLKLTAFVSAASHHPLELHLTQDNLSQFAADLGIPFEFNAINLDAFDPMELIAPTADEVVAVSLPVGCSARTPLPAMLQLVKQLAPKIVVAIDYGSDRSDLPFSQHFLNCLQSCLCLLESLDAAGTDADAVSKIERFLIQPRVEDAVLGRRRADKAIAWRTVLTSAGFAPQPLSNLAEAQADCLLKRVQVRGFHVEKRGAGLALYWQRGELVSVSAWR | LCR1 | 52-70 | GSPSPPNSTSTLSSSHGSG |
| | | | | LCR2 | 79-92 | VAAVSESSAAAAEA |
| | | | | LCR3 | 98-114 | PGEHGGGGGGELPPIPG |
| | | | | LCR4 | 185-215 | SSPAALASDLSSSGGRSLTSSSGSNSKATSA |
| | | | | LCR5 | 221-234 | PEAALQPPPATTAP |
| | | | | LCR6 | 243-255 | PPLLGLPSPTLLL |
| | | | | LCR7 | 277-286 | QQQPLLQPPP |
| | | | | LCR8 | 302-312 | QPQPPPPAPAQ |
| OsGRAS22 | GRAS | 261-634 | LTTLLIHCAQAAAIDDHRNSNELLKQIRQRSSAYGDAGQRLAHCFANALEARLAGTGSNIYRSLAAKRTSVYDILNAFKLYVTACPFKKISNFFSIEAILNASKGMTRLHIVDYGIQYGFQWPIFFQRISKRPGGPPSVRITGVDLPQPGFRPAQLIEATGRRLHDYARMFNVPFEYHAIAAKWDTIRVEDLKIDKDKDELLVVNCLFRMRNMMDEMVTDDSPRMQVLKTIRKMNPNLFIHGVVNGTYNAPFFVTRFKEALFYYSSLFDMLETTASRVDENRLLIERDLFGREALNVVACEGTERVERPETYKQWQVRNIRAGFKQLPLNQETVKKARYKVKKSYHRDFLVDEDNKWMLQGWKGRIIFALSAWE | LCR1 | 2-15 | LDSGSYDDVDYGDL |
| | | | | LCR2 | 56-76 | STPSPTSTTTELENSEDLSES |
| | | | | LCR3 | 237-251 | KGSGNKRGRKKGKSG |
| OsGRAS23 | GRAS | 92-506 | IAAFLADGTCQMQVNDGLSCVVDLAGGDADGGGVGEGRSAQRLASAFAEALALRFILPCDGVCRSLHLTRAPPPPAVSAARQGFRAMCPFVRLAAAAANLSIAEVMEAERAVVHVVDLGGGVDANQWVELVRLVAARPGGPPGLLRLTVVNESEDFLSAVAAYVAAEAQRLDLSLQFHPVLSSIEELSATATGSIGSRLVVIPGQPLAVVANLQIHRLLAFPDYVDGVASRRPAAEQSGSSQHTMTTATKTKADALLRAIRDLNPKLVVLTENEADHNVAELGARVWNALNYYAALFDALEASSTPPAAVPPHERACVERWVLGEEIKDIVVREGTGRRERHETLGRWAERMVAAGFSPVTAARALASTETLAQQMVAAGGGGAGAGVLRAAHGGGCFPVICWCDVPVFSVSTWT | LCR1 | 6-27 | PRLALGGGGGGAGGERLPAAGE |
| | | | | LCR2 | 51-59 | AAMAAAAAA |
| | | | | LCR3 | 513-541 | PAPPLWPPAAAGGAGPSGSGYGGDGPSTA |
| OsGRAS24 | GRAS | 154-558 | LHGHLRRCAEALAASRPADADAELASIARMASSDGDAVQRVAAAFAEAMARVVIRPWRGVSAALFPSDAGAAGDALTAWEAEFARQSFLNLCPLLHLAAVAVNEIILETTRNDKFIHIVDLGGIHHAHWVELLQGLATRRAAVRPCLRLTIVHEHKHFLGQAAQVLAAESDRHGVPLDLHIVESSVEALKLDALGVRSDHAVVIVSTLQLHRLVGAGILSTTAPPSPAAAAAASMITSPLPPANMSSKVDRLLRGFHLLSPRAIILTENEANHFVPSFTDRFASALPYYEQLFAAMEEAGAATVERKAAERYLLREEIKDVIACDHDGPRWARHETLGRWVVRMGAAGFALAPAITVVTAAGRVRAVAARLPGGGDERRYGVTEGGGWLILNREEKPMFCVSAWR | LCR1 | 5-20 | AATAAATTTAAATTAA |
| | | | | LCR2 | 32-52 | MVPVPVASMATATAPAAVAAA |
| | | | | LCR3 | 55-83 | GGHGSSSASQNASGSGEGQGGSMSLSLQL |
| | | | | LCR4 | 89-141 | TPTAAVAVSVPPMAAAPMMAGPAAAAPAPAPPLATMAVAQNASLAAVASALAA |
| OsGRAS25 | GRAS | 116-484 | MIALLMECAAAMSVGNLAGANGALLELSQMASPYAASCGERLVAYFARAMAARLVGSWVGVVAPMAPPPSCGAINAAFRALYNVAPFARLAYLACNQAILEAFHGKRLVHIVDLDVVPGGALQWLSLLPALAARPGGPPVIRVTGFGMSASVLHDTGNQLAGLARKLCMFFEFYAVAKRPGDADAVADMPGRRPGEAVAVHWLRHAMYDAAGDDGASMRLVRWLEPAAVTLVEQERAHGGGGGHGRFLDRFVSALHHYSAVFDAMGASRPDGEDASRHLAEHGVLGREIANVLAVGGPARSSGREGPGSWREVLARHGFAHAGGGGGGRAQLVAAACPGGLGYTVAGDHDGTVRLGWKGTPLYAVSAWT | LCR1 | 13-36 | HHQYLYSSSSSNLPLQQPLLSHHH |
| | | | | LCR2 | 98-111 | ADVEQVAVEDEEEA |
| OsGRAS26 | GRAS | 36-423 | IQQLLLHCAAALESNDVTLAQQAMWVLNNIASSQGDPSQRLTSWLLRALVARACRLCAAAPAGAAVEFLERGRAPPWGRAMSVTELADYVDLTPWHRFGFTASNAAILRAVAGASAVHVVDLSVTHCMQWPTLIDVLSKRPGGAPAIRITVPSVRPAVPPLLAVSSSELGARLAIFAKSKGVQLEFNVVESATTTSPKKTSTTLCQELASVLSDPPSLGLRDGEAVVVNCQSWLRHVAPDTRDLFLDTVRALNPCLLTVTDEDADLGSPSLASRMAGCFDFHWILLDALDMSAPKDSPRRLEQEAAVGRKIESVIGEEDGAERSEPGARLAERMSRKGFAGVVFDEEAAAEVRRLLSEHATGWGVKREDDMLVLTWKGHAAVFTGAWT | LCR | 9-20 | GGGVGAAAHGHG |
| OsGRAS28 | GRAS | 126-479 | LVDDLLDAARLLDAGDSTSAREILARLNHRLPSLPSPPGHAHPPLLRAAALLRDALLPPTALPVSSTPLDVPLKLAAHKALADASPTVQFTTFTSTQAFLDALGSARRLHLLDFDVGFGAHWPPLMQELAHHWRRAAGPPPNLKVTALVSPGSSHPLELHLTNESLTRFAAELGIPFEFTALVFDPLSSASPPLGLSAAPDEAVAVHLTAGSGAFSPAPAHLRVVKELRPAVVVCVDHGCERGALNLLQSCAALLESLDAAGASPDVVSKVEQFVLRPRVERLAVGGGDKLPPPLQSMLASAGFAALQVSNAAEAQAECLLRRTASHGFHVEKRQAALALWWQRSELVSVSAWR | LCR | 35-66 | SSPSTSLGSCSSKPPEDPPPPIAADDDCDWDA |
| OsCIGR1 | GRAS | 201-571 | VKQLLTRCAEALSEDRTEEFHKLVQEARGVVSINGEPIQRLGAYLLEGLVARHGNSGTNIYRALKCREPESKELLSYMRILYNICPYFKFGYMAANGAIAEALRTENNIHIIDFQIAQGTQWITLIQALAARPGGPPRVRITGIDDPVSEYARGEGLDIVGKMLKSMSEEFKIPLEFTPLSVYATQVTKEMLEIRPGEALSVNFTLQLHHTPDESVDVNNPRDGLLRMVKGLSPKVTTLVEQESHTNTTPFLMRFGETMEYYSAMFESIDANLPRDNKERISVEQHCLAKDIVNIIACEGKDRVERHELLGKWKSRLTMAGFRPYPLSSYVNSVIRKLLACYSDKYTLDEKDGAMLLGWRSRKLISASAWH | | | |
| OsGRAS32 | GRAS | 68-434 | LLSLLLRCAEAVAMDQLPEARDLLPEIAELASPFGSSPERVAAYFGDALCARVLSSYLGAYSPLALRPLAAAQSRRISGAFQAYNALSPLVKFSHFTANQAIFQALDGEDRVHVIDLDIMQGLQWPGLFHILASRPTKPRSLRITGLGASLDVLEATGRRLADFAASLGLPFEFRPIEGKIGHVADAAALLGPRHHGEATVVHWMHHCLYDVTGSDAGTVRLLKSLRPKLITIVEQDLGHSGDFLGRFVEALHYYSALFDALGDGAGAAEEEAAERHAVERQLLGAEIRNIVAVGGPKRTGEVRVERWGDELRRAGFRPVTLAGSPAAQARLLLGMYPWKGYTLVEEDGCLKLGWKDLSLLTASSWE | LCR1 | 8-45 | RAPGADAAAMKAKRAADDEEEGGERERARGKRLAAEGK |
| | | | | LCR2 | 55-62 | EEEEAAAE |
| | | | | LCR3 | 437-449 | DGDADADVAVAGD |
| OsCIGR2 | GRAS | 174-544 | LKELLIACARAVEEKNSFAIDMMIPELRKIVSVSGEPLERLGAYMVEGLVARLASSGISIYKALKCKEPKSSDLLSYMHFLYEACPYFKFGYMSANGAIAEAVKGEDRIHIIDFHISQGAQWISLLQALAARPGGPPTVRITGIDDSVSAYARGGGLELVGRRLSHIASLCKVPFEFHPLAISGSKVEAAHLGVIPGEALAVNFTLELHHIPDESVSTANHRDRLLRMVKSLSPKVLTLVEMESNTNTAPFPQRFAETLDYYTAIFESIDLTLPRDDRERINMEQHCLAREIVNLIACEGEERAERYEPFGKWKARLTMAGFRPSPLSSLVNATIRTLLQSYSDNYKLAERDGALYLGWKSRPLVVSSAWH | | | |
| OsSHR1 | GRAS | 192-601 | ASQLLLECARSVAARDSQRVQQLMWMLNELASPYGDVEQKLASYFLQGLFARLTASGPRTLRTLAAASDRNTSFDSTRRTALRFQELSPWSSFGHVAANGAILESFLEVAAAASSETQRFHILDLSNTFCTQWPTLLEALATRSADETPHLSITTVVSAAPSAPTAAVQRVMREIGQRMEKFARLMGVPFRFRAVHHSGDLAELDLDALDLREGGATTALAVNCVNSLRGVVPGRARRRDAFAASLRRLDPRVVTVVEEEADLVASDPDASSATEEGGDTEAAFLKVFGEGLRFFSAYMDSLEESFPKTSNERLALERGAGRAIVDLVSCPASESMERRETAASWARRMRSAGFSPVAFSEDVADDVRSLLRRYREGWSMREAGTDDSAAGAGVFLAWKEQPLVWASAWR | LCR1 | 11-40 | QAASEQQQQQQSASYNSRSTTSSGSRSSS |
| | | | | LCR2 | 46-80 | SYSYYHHSSNSGGGGGGGGYYYGGQQPPPSQYYY |
| | | | | LCR3 | 122-147 | PPASSTPTGTAPTPPLSTSSTAAGAG |
| OsGRAS35 | GRAS | 76-471 | MEQLLVHCANAIEANDATLTQQILWVLNNIAPADGDSNQRLTAAFLCALVSRASRTGACKAVTAAVADAVESAALHVHRFTAVELASFIDLTPWHRFGYTAANAAIVEAVEGFPVVHIVDLSTTHCMQIPTLIDMLAGRAEGPPILRLTVADVAPSAPPPALDMPYEELGAKLVNFARSRNMSMDFRVVPTSPADALTSLVDQLRVQQLVSDGGEALVVNCHMLLHTVPDETAGSVSLTTAQPPVSLRTMLLKSLRALDPTLVVVVDEDADFTAGDVVGRLRAAFNFLWIPYDAVDTFLPKGSEQRRWYEAEVGWKVENVLAQEGVERVERQEDRTRWGQRMRAAGFRAAAFGEEAAGEVKAMLNDHAAGWGMKREDDDLVLTWKGHNVVFASAWA | LCR1 | 9-25 | PPPPPLHPNGHGLGLGL |
| | | | | LCR2 | 34-54 | GGGGARPWSSSSSTTTLGGSG |
| OsSCR1 | GRAS | 283-644 | LLTLLLQCAESVNADNLDEAHRALLEIAELATPFGTSTQRVAAYFAEAMSARLVSSCLGLYAPLPNPSPAAARLHGRVAAAFQVFNGISPFVKFSHFTANQAIQEAFEREERVHIIDLDIMQGLQWPGLFHILASRPGGPPRVRLTGLGASMEALEATGKRLSDFADTLGLPFEFCPVADKAGNLDPEKLGVTRREAVAVHWLRHSLYDVTGSDSNTLWLIQRLAPKVVTMVEQDLSHSGSFLARFVEAIHYYSALFDSLDASYSEDSPERHVVEQQLLSREIRNVLAVGGPARTGDVKFGSWREKLAQSGFRVSSLAGSAAAQAVLLLGMFPSDGYTLIEENGALKLGWKDLCLLTASAWR | LCR1 | 3-55 | SSSLLLFPSSSSSATHSSYSPSSSSHAITSLLPPLPSDHHLLLYLDHQEQHHL |
| | | | | LCR2 | 86-101 | AAAAPSSASAQLPALP |
| | | | | LCR3 | 115-134 | AAPAPPPPQQQVAAGEGGPP |
| | | | | LCR4 | 148-157 | ASSGAAVSVA |
| | | | | LCR5 | 188-229 | SDPAPPPPPPPSHPALLPPDATAPPPPPTSVAALPPPPPPQP |
| | | | | LCR6 | 235-279 | EPQCQEQEPNQPQSPKPPTAEETAAAAAAAKERKEEQRRKQRDEE |

| | | | | | | |
|---|---|---|---|---|---|---|
| ΨOsGRAS8 | GRAS | 199-356 | LRELLMSCAQAVASGNRRSAGELLEQIKRHSSPTGDATERLAHYFADGLEARLAGAASLERRLVASAEERASAMELLEAYQVFMAACCFKWVAFTFANMAILRAAEGRNRLHIVDYGGQYHGLQWPSLLQRLAEREGGPPEFRAVAAARWETVTAEDV | LCR1 | 29-40 | PAAPPSEAAAAA |
| | GRAS | 339-542 | EFRAVAAARWETVTAEDVVGVDPDDEAAVVVNDVLSLGTLMDESGVFDDPSPRDTVLGSIRDMRPAVFVQAVVNGAHGAPFFPTRFREALFFFSALFDMLGATTPEEGSHLRVVLERDVLRRAAVGVIAGEGAERVERPETYRRWQARNRRAGLRQAAVEGDVVEAVRRRVRRRHHEEFVIEEDAGWLLQGWKGRILYAHSAWV | LCR2 | 108-122 | GSGNGRGRKGSKHGG |
| | | | | LCR3 | 140-147 | EEEEDDDD |
| | | | | LCR4 | 177-193 | AEKKCGKAARRRRRQAK |
| OsGRAS39 | GRAS | 67-365 | RDVLVVHIVDLSCSAAHPWQWPKLLDDFHGRPGGAPELYLTVLHDDNDFLADMQSLLSKKAESLGVSFHFISVIGRLETLDFSNLRSTFQIKFGVAVAISCALQMHRLLLVDDNLSSTSIAQLQKMANFTQPKQMASSVCSPASTLNYLQTPSPRTPKLLARLLSAIRALKPNIMLIMEQDADHNTLLFRDRFNEVLNYYAALFDCFHAVAAANPGRTDERLRVDRMILREEIKNILVCEGVHRHERHERLDQWAMHMEESGFHNVQLSFSAIREAYVWQLKVQADNLRLCCTDRGMFQ | LCR | 370-391 | SSATSSPASSVYSPSPSPSNGS |
| | GRAS | 405-843 | LIGLLYQCAAEVSAGSFDRANLCLEHITQLASLDAPHALQRLAAVFADALARKLLNLILGLSRALLSSANSADAHLVPVARRHMFDVLPFLKLAYLTTNHAILEAMEGERFVHVVDFSGPAANPVQWIALFHAFRGRREGPPHLRITAVHDSKEFLANMAAVLSKEAEAFDIAFQFNAVEAKLDEMDFDALRHDLGVRSGEALAVSVVLQLHRLLAVDDGRRHAAAGCLTPVQIIARSSPRSFGELLERELNTRLQLSPDASVVSSLSPHSPAAATAAHPTTSTPKLGSFLSAVRSLSPKIMVMTEQEANHNGGAFQERFDEALNYYASLFDCLQRSAAAAAERARVERVLLGEEIRGVVACEGAERVERHERARQWAARMEAAGMERVGLSYSGAMEARKLLQSCGWAGPYEVRHDAGGHGFFFCWHKRPLYAVTAWR | | | |
| OsGRAS41 | GRAS | 4-339 | LSDLLLAGAEAVEAGDSILASVAFSRLDDFLSGIPENGAASSFDRLAYHFDQGLRSRMSSASTGCYQPEPLPSGNMLVHQIIQELSPFVKFAHFTTNQAILDAIIGDMDVHVVDLNIGEGIQWSSLMSDLARCGGKSFRLTAITTYADCHASTHDTVVRLLSEFADSLELPFQYNSICVHNEDELHAFFEDCKGSVIVSCDTTSMYYKSLSTLQSLLLVCVKKLQPKLVVTIEEDLVRIGRGVSPSSASFVEFFFEALHHFTTVFESMASCFIGSSYEPCLRLVEMELLGPRIQDFVVKYGSVRVEANASEVLEGFMACELSACNIAQARMLVGLFNRVFGVVFKKISLLMVY | | | |
| | Transmemebrane region | 354-376 | VIWSSLAAGCGSHGIVVLAFYAA | | | |
| ΨOsGRAS9 | GRAS | 1-129 | MSHLENTLEARLAGTGSQMYQSLVAKRTSTVDFLKAYKLFTAACCVKKTIYNAVAGKRKLHIVDYGLSYGFQWPALFFLLGTREGGPPEVRMTGIDVPQPGFRPADQIEETGRRLSICARAPVRCAIQV | LCR | 216-254 | SRSAPSSSPPRPSLHLHLHLRRRPPPSSSRHAADDAALH |
| OsGRAS43 | RPT1 | 94-113 | ASPRRDFMACSPKRDYMVTT | LCR1 | 2-13 | AGGGAKLQQQQA |
| | RPT1 | 114-134 | SSPKRDYMVTSSPKRDYMVSS | LCR2 | 165-177 | HGGGGGGGHHLHH |
| | GRAS | 401-771 | LVHLLLACADLVSKGDHPAALRHLHLLRRVASPLGDSMQRVASHFADALAARLSLLSSPTSASPSPRAAAAAAPYPFPPSPETLKVYQILYQACPYIKFAHFTANQAIFEAFHGEDRVHVVDLDILQGYQWPAFLQALAARPGGPPTLRLTGVGHPPAAVRETGRHLASLAASLRVPFEFHAAAADRLERLRPAALHRRVGEALAVNAVNRLHRVPSSHLPPLLSMIRDQAPKIITLVEQEAAHNGPYFLGRFLEALHYYSAIFDSLDATFPAESTARMKVEQCLLAPEIRNVVACEGAERVARHERLERWRRLMEGRGFEAVPLSAAAVGQSQVLLGLYGAGDGYRLTEDSGCLLLGWQDRAIIAASAWR | LCR3 | 183-201 | GGGMEGGGGGHGAQPQYGG |
| | | | | LCR4 | 223-230 | GGGSGGGG |
| | | | | LCR5 | 280-288 | GGVGGGGGG |
| | | | | LCR6 | 293-306 | SGASVSVVTAPASS |
| | | | | LCR7 | 326-341 | GGGDEAVAAAMAVAGE |
| | | | | LCR8 | 355-364 | GGGGEFGGEG |
| OsGRAS44 | GRAS | 303-683 | LHTLLIHCAQAVATSDRRSATELLKQIKQNSSARGDATQRLACCFAEGLEARLAGTGSQVYKSLVAKCTSTVDFLKAYKLFAAACCIKKVSFIFSNKTILDAVAGKRKLHIVDYGLSYGFQWPGLFKCLSEREGGPPEVRITGIDFPQPGFRPADQIEETGRRLSNCARQFGVPFRFQAIAAKWETVRREDLHLDREEEEEEEEEVLVVNCLHFLNALQDESVVVDSPSPRDMVLNNIRDMRPHVFVQCVVNGAYGAPFFLTRFRETLFFYSSQFDMLDATIPRDNDERLLIERDILGRWALNVIACEGADRVDRPETYKQWLVRNHRAGLTQLPLQPQVVELVRDKVKKLYHKDFVIDVDHNWLLQGWKGRILYAMSTWV | LCR1 | 14-27 | LEPFSPSLFLDLPP |
| | | | | LCR2 | 82-111 | SDDTTTNSSDDSASATTNNTTNSAAAANAS |
| | | | | LCR3 | 209-218 | GRSGGSGRGR |
| | | | | LCR4 | 279-299 | AEKKARNGGGAGRRAARAKAA |
| OsGRAS47 | GRAS | 215-586 | LRMLLIQCAQAMATDNQQSAGELLKKIKQHALATGDAMQRVAHYFAKGLEARLAGSGKHLYQNHVRMSLVEYLKVYKLYMAACCFKKVALMFAAMTIMQAVQGKKRLHIVDYGIRCGLHWPDLFRRLGSREDGPPEVRITIVDIPQPGFRPFQRIEAAGHCLSSCANEFRVPFRFQAVVAAKWETVGAEDLHIEPDEVLVVNDLWSFSALMDESIFCDGPNPRDVALRNISKMQPDVFIQGIINGGYGASFLSRFRGALLYYSALFDMLDATTPRESGLRLALEQNVLGPYALNAIACEGADLVERPEKYRQWQARNHRAGMQQLKLRPDIVDTIREEVNKYHHKDFLLGEDGQWLLQGWMGRVLFAHSAWV | LCR | 201-211 | KKKGKKGSSSK |
| ΨOsGRAS10 | RPT1 | 61-73 | FLDMMVIQESANE | LCR1 | 22-30 | SSSSLLLWS |
| | RPT1 | 117-129 | FLEMMAIQESAND | LCR2 | 47-60 | DADHSHDQIHQDHQ |
| | GRAS | 173-308 | AGDLLLAGAMAVDAGDAVHASAIMSRLDDLLADIAGRRSCEATSPVDHLAYYFARGLKLRISGAATPASSPPPPAANWSSPAYRMLQELTPFVKFAHFTANQAILEATADDLDVHVVDFNVGEGVQWSSLMLKLLL | | | |
| | GRAS | 305-460 | KLLLLGTITILQPKLVILIEDELSRISKNPPSPSLAAPPPFPEFFSDAVAHFTAVMESTASCLVSYDDEAWLSLRRVGEEVVGPRVEDAVGRYGSLAGGAQMMEGLRAREVSGFSVAQGKMLAGLFGGGFGVVHQEKGRLALCWKSRPLISVSLWC | | | |
| OsGRAS53 | RPT1 | 190-207 | FLKGMEEANKFLPTENKL | LCR1 | 53-65 | PPSPPPPTTTATT |
| | RPT1 | 219-236 | YLRGLEEAKRFLPSDDKL | LCR2 | 116-135 | LSDPSSNSRSSNSDDPRLSP |
| | GRAS | 358-730 | LRTLLIHCAQAVATDDRRSATELLKQIKQHAKPTGDATQRLAHCFAEGLQARIAGTGSLVHQSLVAKRTSAVDILQAYQLYMAAICFKKVSFIFSNQTIYNASLGKKKIHIVDYGIQYGFQWPCFLRRISQREGGPPEVRMTGIDLPQPGFRPTERIEETGHRLSKYAQEFGVPFKYNAIAAVKMESVRKEDLNIDPDEVLIVNCQYQFKNLMDESVVIDSPRDIVLSNIRKMQPHVFIHAIVNGSFSAPFFVTRFREALFFYSALFDVLDATTPRESEQRLLIEQNIFGRAALNVIACEGIDRVERPETYKQWQVRNQRAGFKQLPLNPEIVQVVRNKVKDCYHKDFVIDIDHQWLLQGWKGRILYAISTWT | LCR3 | 150-162 | AAATATAVAAAAV |
| | | | | LCR4 | 241-260 | AAAAAPVVSVKKEAVDVVVA |
| | | | | LCR5 | 337-356 | GGKGGNGKVKGGRRGGRDVV |
